# YTHDF2 upregulation and relocation dictate CD8 T cell polyfunctionality in tumor immunity

**DOI:** 10.1101/2024.07.11.603088

**Authors:** Haiyan Zhang, Xiaojing Luo, Wei Yang, Zhiying Wu, Zhicong Zhao, Xin Pei, Xue Zhang, Chonghao Chen, Josh Haipeng Lei, Qingxia Shi, Qi Zhao, Yanxing Chen, Wenwei Wu, Zhaolei Zeng, Huai-Qiang Ju, Miaozhen Qiu, Jun Liu, Bin Shen, Minshan Chen, Jianjun Chen, Chu-Xia Deng, Rui-Hua Xu, Jiajie Hou

## Abstract

Epigenetic traits impact the antitumor function of CD8 T cells, yet whether and how RNA methylation programs engage in T cell immunity is poorly understood. Here we show that the *N*^6^-methyladenosine (m^6^A) RNA reader YTHDF2 is highly expressed in early effector or effector-like CD8 T cells and is partially distributed in the nucleus. YTHDF2 loss in T cells exacerbates tumor progression and confers unresponsiveness to PD-1 blockade in mice and humans. In addition to initiating RNA decay for mitochondrial fitness, YTHDF2 can orchestrate chromatin regulation to promote T cell polyfunctionality. YTHDF2-mediatd preservation of gene transcription arises from the interaction of YTHDF2 with IKZF1/3. Accordingly, immunotherapy-induced efficacy could be largely restored in YTHDF2-deficient T cells through combinational use of lenalidomide. Moreover, m^6^A recognition is fundamental for YTHDF2 translocation to the nucleus and autoregulation at the RNA level. Thus, YTHDF2 coordinates epitranscriptional and transcriptional networks to potentiate T cell immunity.

**Highlights:**

- YTHDF2 expression and distribution underpin the threshold for bona fide CD8 T cell effector response
- Canonical YTHDF2-mRNA decay pathway alleviates mitochondrial stress and CD8 T cell exhaustion
- Nuclear YTHDF2 sequesters IKZF1/3-mediated transcriptional repression to safeguard CD8 T cell polyfunctionality
- The tumoricidal activity of YTHDF2-deficient CD8 T cells could be repaired through the synergy of anti-PD-1 and lenalidomide

## Introduction

Among more than 170 types of RNA modifications, *N^6^*-methyladenosine (m^6^A) represents the most prevalent and abundant modification in eukaryotic mRNA. By enlisting the “writer” (methyltransferase), “eraser” (demethylase) and “reader” proteins, dynamic m^6^A modification regulates nearly every step of mRNA metabolism and interferes with various biological processes^1^. Emerging evidence shows that m^6^A RNA modifiers within tumor cells or surrounding myeloid cells largely affect tumor immunity and immunotherapy efficacy^2,3^. For instance, the tumor-intrinsic m^6^A demethylase FTO can either subvert the host immune reaction by facilitating the expression of the immune checkpoint gene *LILRB4*^1^ or restrict T cell activation by altering the metabolic microenvironment^4^. The methyltransferase complex component METTL14 disrupts the interferon-γ signalling within microsatellite instability-low tumors and limits the response to immune checkpoint blockade (ICB) therapy^5^; however, METTL14 governs m^6^A RNA stabilization in a subpopulation of tumor-associated macrophages to prevent T cell dysfunction^6^. The m^6^A reader YTH domain family 1 (YTHDF1) in classic dendritic cells impedes the cross-presentation of tumor antigens and the cross-priming of CD8 T cells^7^. Two recent reports showed that myeloid cell YTHDF2 is also associated with tumor immunosuppression^8,9^. These findings indicate that the m^6^A machinery posttranscriptionally controls cancer-immune set points, which may be useful targets for overcoming immunotherapy resistance.

Tumor-reactive CD8 T cells are key to both natural and therapy-induced antitumor immunity. However, chronic exposure to tumor antigens or environmental stimuli may render CD8 T cells dysfunctional and limit the outcomes of cancer therapy^10,11^. Remarkably, T cell activation and differentiation are often concomitant with epigenetic processes, many of which account for the molecular rewiring of T cell dysfunction imposed by the tumor microenvironment (TME)^12^. Therefore, manipulating T cell epigenetic programs may foster cancer therapeutic efficacy owing to the acquisition of long-term T cell persistence^13,14^. Although epigenetic changes accompanying differentiation have been extensively studied^15^, how T cell effector polyfunctionality is epigenetically safeguarded in the early phase remains elusive. In addition, recent insights have revealed that a progenitor exhausted T (T_pex_) cell population can partially differentiate into effector-like transitory exhausted T (T_ex_) cells, which serve as a cardinal force of T cell immunity when responding to anti-PD-1 therapy^16,17^, highlighting the need for knowledge assimilation to better understand and harness this functional process. Given both distinct and shared epigenetic circuits between different T cell subsets^18^, we embark on identifying a novel regulator that governs early epigenetic events exclusively for tumoricidal effector and effector-like T cells. m^6^A-mediated RNA methylation and destabilization have been recognized as an ingenious mechanism for T cell homeostasis^19^ and survival^20^, but the implication of m^6^A RNA modifiers in antitumor T cells remains an enigma. Therefore, it is imperative to illustrate the m^6^A machinery underlying T cell activation and therapy-induced rejuvenation.

RNA m^6^A is cotranscriptionally installed by the methyltransferase complex in the nucleus^21^. Recent studies have suggested that m^6^A modification has an interplay with and an impact on the chromatin state. In mouse embryonic stem cells, METTL14 can recognize transcription elongation mark histone H3 trimethylation at Lys36 (H3K36me3), which guides m^6^A deposition on actively transcribed nascent RNAs^22^. Conversely, m^6^A-modified nuclear RNAs can direct chromatin organization and gene expression. The nuclear m^6^A reader YTHDC1 plays a sophisticated role in such a context; it can either recruit the histone demethylase KDM3B to erase the repressive histone mark H3K9me2, or dictate nuclear RNA decay to restrict the chromatin activity and downstream transcription ^23–25^. These discoveries raise questions about whether the m^6^A machinery adapts the binding interface for core transcription factors and whether T cell activation necessitates the crosstalk between m^6^A modification and chromatin organization.

YTHDF2, a highly effective m^6^A reader, specifically recognizes and degrades m^6^A-containing mRNAs in the cytoplasm, where it primarily resides^26^. Otherwise, under heat shock stress, nucleus-localized YTHDF2 protects the 5’ untranslated region (5’UTR) of stress-induced transcripts from FTO-mediated demethylation, resulting in cap-independent translation initiation^27^. In the present study, we reveal that YTHDF2 is uniquely expressed and distributed during early T cell activation and therapy-induced rejuvenation. Despite its short-term upregulation and nuclear localization, YTHDF2 ensures the longevity and tumoricidal activity of CD8 T effector (T_eff_) and T_eff_-like cells. In quiescent T cells, cytoplasmic YTHDF2 potentially destabilizes its cognate coding mRNA via m^6^A recognition, self-maintaining a low expression level under nonpathological conditions. When encountering robust antigen stimuli or ICB therapy, a portion of YTHDF2 switches to a non-autoregulating state followed by nuclear translocation, allowing cognate mRNA translation in early polyfunctional T cells. The accumulation of YTHDF2 in the cytoplasm results in the degradation of the redundant mitochondrial component-encoding mRNAs, withstanding mitochondrial stress and T cell exhaustion. In a more tumor-rejecting manner, the nuclear import of YTHDF2 directs chromatin organization and effector polyfunctionality by limiting IKZF1 and IKZF3 from the transcriptional inhibition of T cell receptor (TCR) signalling. Conversely, YTHDF2 deficiency in T cells thwarts both endogenous and ICB-induced tumor immunity in mice and is correlated with a poor therapeutic response in cancer patients. Nevertheless, owing to the dominant transcriptional repression by IKZF1/3 in YTHDF2-null T cells, lenalidomide, a clinically available IKZF1/3 inhibitor, could achieve a compensatory immune response in synergy with ICB. Collectively, these data provide proof-of-concept evidence that YTHDF2 integrates RNA and DNA epigenetics to potentiate T cell antitumor immunity.

## Results

### YTHDF2 is selectively upregulated and redistributed in early T_eff_ and T_eff_-like cells

To begin, we interrogated several transcriptomic datasets and assessed the expression of m^6^A machinery components in T cells that undergo diverse signals. When stimulated with anti-CD3/CD28 antibodies, both CD4 and CD8 T cells showed a predominant increase in *Ythdf2* expression during early activation (Supplementary Fig. 1a, b). In an *in vitro* system mimicking different states of human CD8 T cells^28^, a high *Ythdf2* mRNA level was observed during 3–48 h of anti-CD3/CD28 stimulation, but its expression decreased at later time points of activation as well as towards exhaustion-like and memory-like stages (Fig. 1a). Similarly, among the *in vitro*-generated CD4 T cell subsets, a group of inducible tolerant T cells exhibited the lowest YTHDF2 level (Supplementary Fig. 1c). In immunotherapy settings, PD-1 blockade induced the upregulation of YTHDF2 in CD8 T cells from responding tumors; however, this upregulation occurred only during early tumor regression but not in the late regression or progression stage (Fig. 1b). As shown in these datasets, the expression of the gene encoding another important m^6^A reader, YTHDF1, could also be upregulated upon T cell activation and rejuvenation, but its expression level was relatively low in tumor-infiltrating CD8 T cells (Supplementary Fig. 1d). A recent report indicated that the loss of YTHDF1 in CD8 T cells does not affect antitumor immunity^7^. Therefore, the potential function of YTHDF2 in T cell-mediated tumor immunity was the focus of this study.

**Fig. 1.**
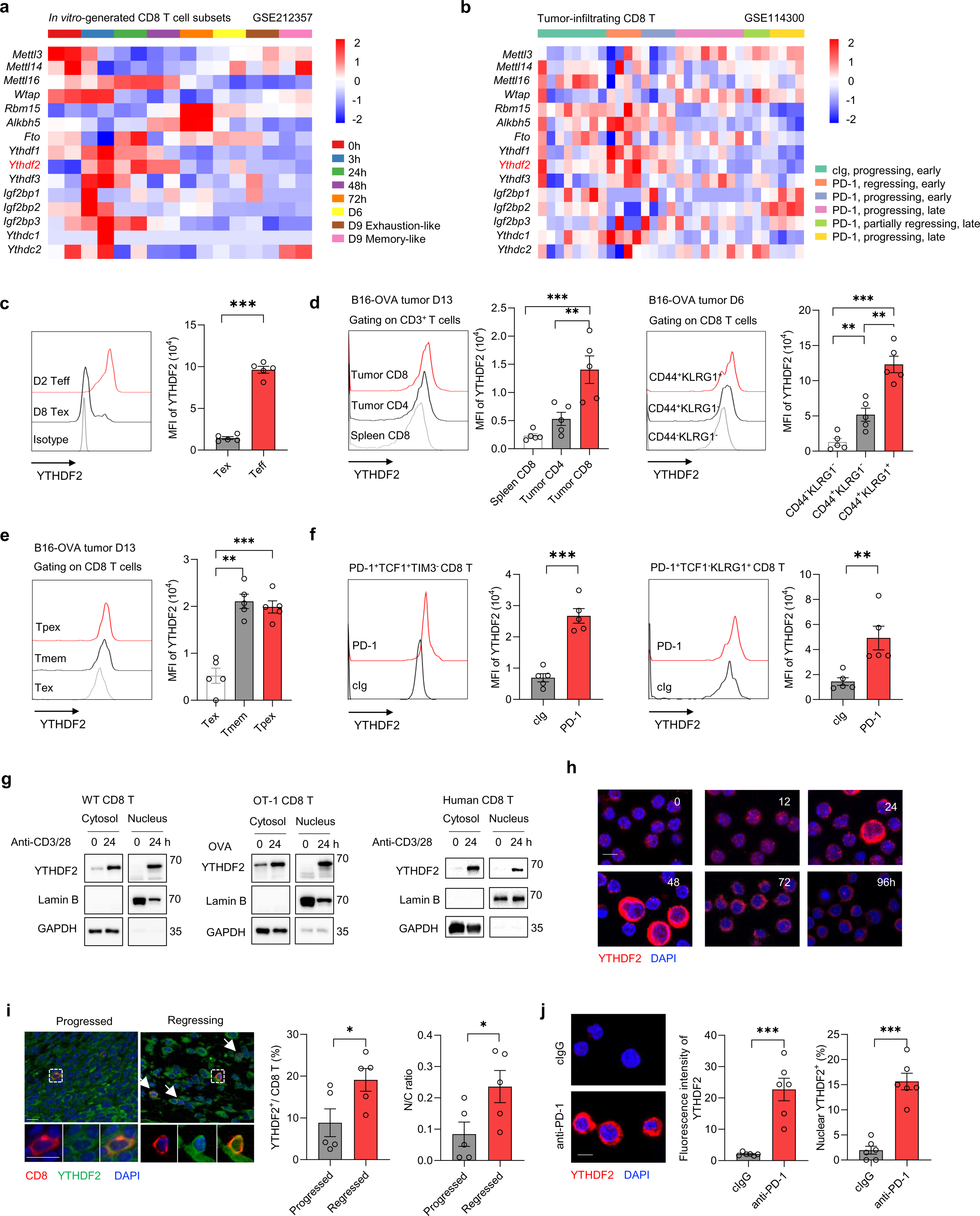
YTHDF2 is selectively upregulated and redistributed in early T_eff_ and T_eff_-like cells. **a–b** Transcriptomic data were mined from existing datasets. mRNA expression of m^6^A modifiers in CD3/CD28 bead-based stimulation of human CD8 T cells at various time points indicating naive/memory, activated, and exhausted populations (GSE212357) (a). mRNA expression of m^6^A modifiers in total tumor-infiltrating CD8 T cells from HKP1 lung cancer-bearing mice upon anti-PD-1 or cIg treatment (GSE114300) (b). **c** Quantification of YTHDF2 MFI between *in vitro*-generated effector CD8 T cells (n = 5) and exhausted CD8 T cells (n = 5). **d–e** Quantification of YTHDF2 MFI in various cell populations from late time point (Day 13) B16F10-OVA tumor (n = 5) and spleen (left panels) or from early time point (D6) B16F10-OVA tumor (n = 5) by flow cytometry. **f** Quantification of YTHDF2 MFI in T_pex_ cells and transitory T_ex_ cells from B16F10-OVA tumor treated with anti-PD-1 (250μg/mouse) (n = 5) or control IgG (cIg) (n = 5). **g** Immunoblotting analysis of YTHDF2 in the cytosol and nucleus of mouse (WT or OT-1) and human CD8 T cells stimulated with anti-CD3/CD28 (5 μg/ml) or OVA (10 nM). **h** Representative confocal immunofluorescence images of YTHDF2 (red) and DAPI (blue) in naïve (n = 5) or activated (anti-CD3/CD28, 5 μg/ml, 12–96h) CD8 T cells (n = 5). Scale bar, 10 μm. **i** Representative immunofluorescent staining of CD8 (red) and YTHDF2 (green) in regressing (n = 5) or progressed (n = 5) B16-OVA tumors. A dashed box represents the 4× enlarged area shown in the bottom panels with separate channels. White arrows point to cells positive for YTHDF2 and CD8. Scale bar, 10 μm. Middle panel, frequencies of YTHDF2-positive CD8 T cells. Right panel, quantification of the nuclear to cytoplasmic ratios of YTHDF2 intensity in YTHDF2-positive CD8 T cells. **j** Representative confocal immunofluorescence images of YTHDF2 (red) and DAPI (blue) in PD-1^+^SLAMF6^+^TIM3^-^ CD8 T_pex_ cells from B16-OVA tumors (Day 16) after *in vitro* anti-PD-1 (10 μg/ml, 48 h) or cIg stimulation (n = 6). Scale bar, 10 μm. Error bars, mean ± s.e.m. \**P* < 0.05; \*\**P* < 0.01; \*\*\**P* < 0.001. One-way analysis of variance (ANOVA) (d, e) or two-tailed unpaired Student’s t-test (c, f, i, j).

We obtained both human peripheral and mouse splenic CD8 T cells for *in vitro* activation and validated the inducible upregulation of YTHDF2 expression at the protein level (Supplementary Fig. 1e, f). By performing liquid chromatography-tandem mass spectrometry (LC–MS/MS) analysis, we observed a significant loss of m^6^A modification in the transcriptome after CD8 T cell activation (Supplementary Fig. 1g). We then conducted antibody-based m^6^A sequencing (m^6^A-seq) and noted that m^6^A-hypomethylated peaks were mostly distributed in 3’ untranslated regions (3’UTRs) (Supplementary Fig. 1h). In addition to the possibility of mRNA 3′UTR shortening pending T cell activation^29^, this observed hypomethylation might also be explained by YTHDF2-induced destabilization of m^6^A-containing mRNAs^26^.

As shown by flow cytometry analysis, *in vitro*-generated early-phase effector CD8 T cells manifested a much higher YTHDF2 level than did exhausted CD8 T cells (Fig. 1c). In C57BL/6 mice subcutaneously challenged with ovalbumin (OVA)-expressing B16F10 cells, tumor-infiltrating T cells expressed more YTHDF2 than did splenic T cells, among which CD44^+^KLRG1^+^ terminal effector CD8 T cells were predominant (Fig. 1d). Consistent with the transcriptomic data, PD-1^+^TIM3^+^TCF1^-^ terminal exhausted CD8 T cells exhibited a much lower YTHDF2 level than did PD-1^+^TCF1^+^SLAMF6^+^TIM3^-^ T_pex_ and CD44^+^PD-1^-^TCF1^+^CD127^+^ memory T cell (T_mem_) subsets (Fig. 1e). In addition, early anti-PD-1 treatment led to a 3-fold increase in YTHDF2 expression in T_pex_ and their progeny PD-1^+^TCF1^-^KLRG1^+^ transitory T_ex_ cells (also known as terminal T_eff_-like cells^30^, which are believed to expand upon ICB for tumor killing^16,17,31^ (Fig. 1f). These observations imply that YTHDF2 may widely impact tumor-experienced CD8 T cells, particularly the early effector and effector-like subsets.

Whereas it is well-accepted that YTHDF2 primarily resides in the cytosol, where mRNA decay occurs, previous work has shown that heat shock stress can lead to the relocation of YTHDF2 to the nucleus through an unknown mechanism^27^. In the present study, we assessed whether T cell activation or reinvigoration could alter the subcellular localization of YTHDF2. Surprisingly, wild-type and OT-1 CD8 T cells accumulated nuclear YTHDF2 when stimulated with anti-CD3/CD28 antibodies and OVA peptides, respectively, for 12–48 h (Fig. 1g, h and Supplementary Fig. 1i, j). In the case of the Jurkat human T lymphoma cell line, we detected an inherent nuclear fraction of YTHDF2, which modestly increased after phytohaemagglutinin (PHA) treatment (Supplementary Fig. 1k, l). Akin to the temporary YTHDF2 relocation observed *in vitro*, tumor-infiltrating CD8 T cells showed nuclear expression of YTHDF2 in regressing but not progressed lesions (Fig. 1i). We then implanted OT-1 (expressing a TCR specific to MHC-I-restricted OVA residues) transgenic mice with B16F10 or B16F10-OVA cells but discovered YTHDF2-redistributed CD8 T cells only within tumors formed by the latter (Supplementary Fig. 1m), showing that such a phenotype depends on antigen-specific T cell reactions. As expected, early anti-PD-1 treatment triggered the overexpression and nuclear relocation of YTHDF2 within a small portion of CD8^+^ tumor-infiltrating lymphocytes (TILs) (Supplementary Fig. 1n). Specifically, PD-1^+^SLAMF6^+^TIM3^-^ CD8 T_pex_ sorted from B16-OVA tumors exhibited YTHDF2 overexpression and nuclear relocation when cultured in the presence of anti-PD-1 (Fig. 1j). Mirroring the selective overexpression pattern, these data highlight subcellular YTHDF2 distribution as an acute T cell phenotype underlying natural or therapy-induced tumor eradication, which prompted us to explore the multiple functions of YTHDF2 in T cell immunity.

### YTHDF2 is essential for the antitumor effects of CD8 T cells

We crossed *Ythdf2^Flox/Flox^* (hereafter *Ythdf2^F/F^*) mice^32^ with *dLck^Cre^* transgenic mice (expressing Cre recombinase under the distal *Lck* promoter)^33^ to conditionally knockout *Ythdf2* in T cells (*Ythdf2^CKO^*) (Supplementary Fig. 2a, b). First, we compared the thymuses, spleens and peripheral blood from 6-week old *Ythdf2^F/F^*, *dLck^Cre^* and *Ythdf2^CKO^* mice and found no obvious differences in their T cell compositions (Supplementary Fig. 2c). Gene knockout efficiency was demonstrated by comparing the intratumoral CD8 T cells from *Ythdf2^F/F^* and *Ythdf2^CKO^* mice (Supplementary Fig. 2d). When inoculated with hepatocellular carcinoma Hepa1-6 cells, melanoma B16F10 cells, or colorectal carcinoma MC38 cells, *Ythdf2^CKO^* mice exhibited much faster tumor growth than did *Ythdf2^F/F^* or *dLck^Cre^* mice (Fig. 2a–c and Supplementary Fig. 2e). Correspondingly, *Ythdf2* deficiency led to lower numbers of tumor-infiltrating CD8 T cells but did not affect CD4 T cells or regulatory T cells (Fig. 2d and Supplementary Fig. 3a, c). Dextramer staining further indicated that the frequency of the tumor-specific CD8 T cell subpopulation was substantially decreased in the absence of YTHDF2 (Supplementary Fig. 3a). Reduced CD8 T cell numbers and percentages were also seen in the tumor-draining lymph nodes (dLNs) of *Ythdf2 ^CKO^* mice (Supplementary Fig. 3b). Moreover, the percentage of terminal effector CD8 T cells was lower in *Ythdf2^CKO^* mice at day 12 after MC38 tumor inoculation, consistent with increased apoptosis and impaired cytokine production and proliferative capacity of a broader CD8 T cell population (Fig. 2e, f). In contrast, PD-1^+^TIM3^+^CD101^+^ terminally exhausted CD8 T cells were more frequently found in the *Ythdf2^CKO^* group at a later time point (Fig. 2g and Supplementary Fig. 3d). In addition to the crucial role of CD8 T cell immunity, we were also curious about whether CD4 T cells were affected in *Ythdf2^CKO^* mice. However, antibody-mediated neutralization confirmed that YTHDF2 loss mainly jeopardizes CD8 (but not CD4) T cell-mediated antitumor immunity (Supplementary Fig. 3e), possibly because YTHDF2 expression by regulatory T cells is conductive to tumor growth^34^.

**Fig. 2.**
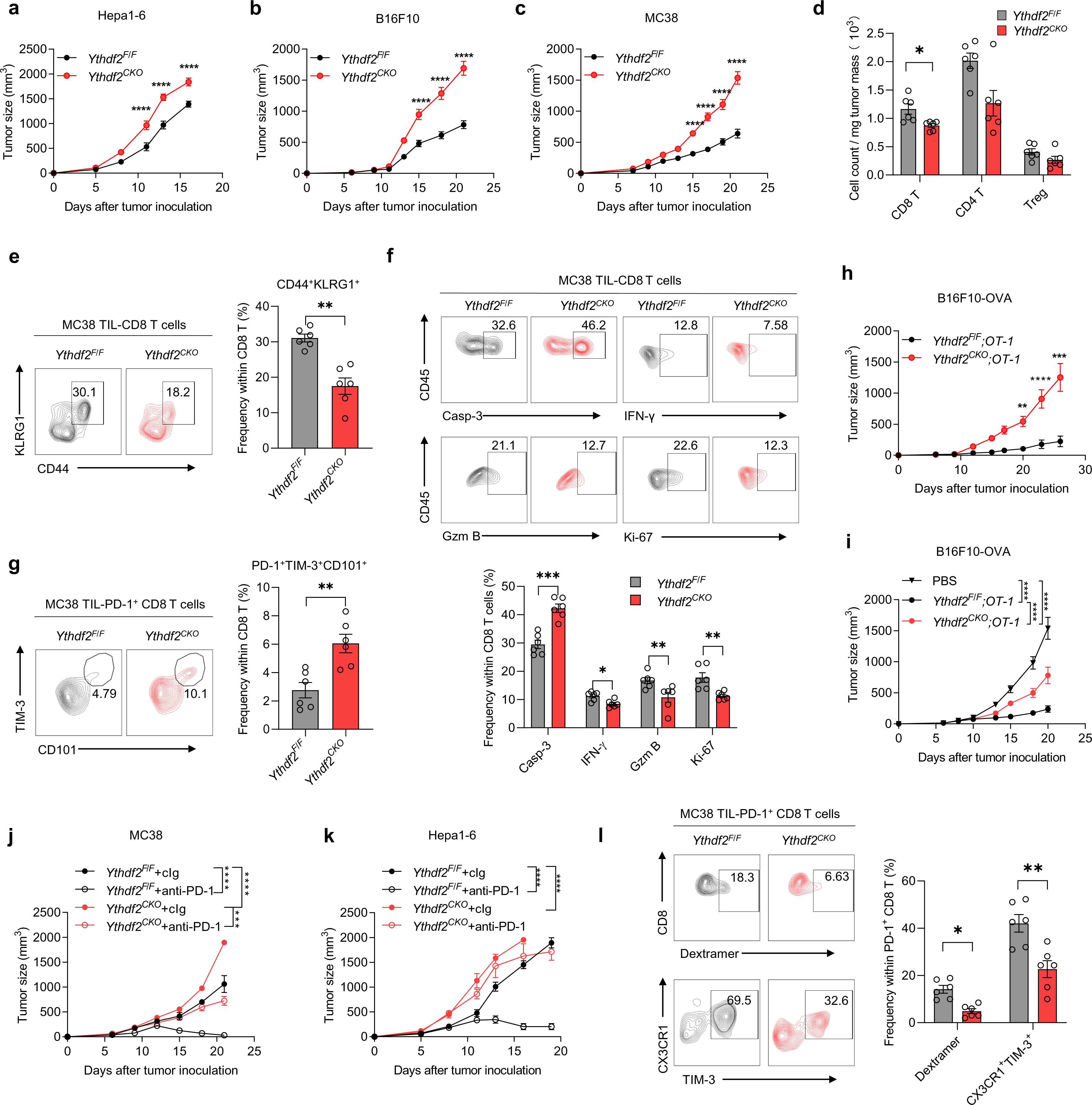
YTHDF2 is essential for the antitumor effects of CD8 T cells. **a** Male *Ythdf2^F/F^* (n = 6) or *Ythdf2^CKO^* (n = 6) mice were injected subcutaneously with 10^6^ Hepa1-6 cells. Tumor growth was monitored ever 2 or 3 days. **b–c** Female*Ythdf2^F/F^* (n = 6) and *Ythdf2^CKO^* (n = 5–6) mice were injected subcutaneously with 5 × 10^5^ B16F10 (b) or 10^6^ MC38 (c) cells. Tumor growth was monitored ever 2 or 3 days. **d–f** Tumor-infiltrating lymphocytes (TILs) were isolated from *Ythdf2^F/F^* (n = 6) and *Ythdf2^CKO^* (n = 6) mice 12 days after MC38 tumor inoculation. Numbers of immune subsets (CD8 T, CD4 T and T_reg_ cells) within TILs (d) and frequencies of CD8 T cell subpopulations positive for CD44^+^KLRG1^+^ (e) active caspase-3 (Casp-3), granzyme B (Gzm B), IFN-γ or Ki-67 (f) were assessed by flow cytometry. **g** Frequencies of PD-1^+^TIM3^+^CD101^+^CD8 T cell subpopulations from MC38 tumor-bearing *Ythdf2^F/F^* (n = 6) and *Ythdf2^CKO^* (n = 6) mice (Day 18). **h** Female*Ythdf2^F/F^;OT-1* (n = 8) or *Ythdf2^CKO^;OT-1* (n = 6) mice were injected subcutaneously with 10^6^ B16F10-OVA cells and monitored for tumor growth. **i** Adoptive transfer therapy using PBS control or OVA-primed (72 h) *Ythdf2^F/F^;OT-1* or *Ythdf2^CKO^;OT-1* CD8 T cells against B16F10-OVA melanoma (n = 5 per group). **j–k** Female *Ythdf2^F/F^* (n = 12) or *Ythdf2^CKO^* (n = 10) mice were injected subcutaneously with 10^6^ MC38 cells (j). Male *Ythdf2^F/F^* (n = 16) or *Ythdf2^CKO^* (n = 12) mice were injected subcutaneously with 10^6^ Hepa1-6 cells (k). Tumor-bearing mice were treated with anti-PD-1 (250 μg/mouse) or cIg. **l** TILs were isolated from *Ythdf2^F/F^* (n = 6) and *Ythdf2^CKO^* (n = 6) mice 12 days after MC38 tumor inoculation with anti-PD-1 treatment. Frequencies of CD8 T cell subpopulations positive for PD-1^+^Dextramer^+^, CX3CR1^+^Tim3^+^CD101^-^PD-1^+^ were assessed by flow cytometry. Error bars, mean ± s.e.m. \**P* < 0.05; \*\**P* < 0.01; \*\*\**P* < 0.001; \*\*\*\**P* < 0.0001. Two-way ANOVA (a–c, h–k) or two-tailed unpaired Student’s t-test (d-g, l).

To determine whether YTHDF2 engages in regulating tumor-specific CD8 T cells, we further bred *Ythdf2^CKO^* (or *Ythdf2^F/F^*) mice with OT-1 transgenic mice to generate a *Ythdf2^CKO^*;OT-1 (or *Ythdf2^F/F^*;OT-1) line. Strikingly, OVA-expressing B16F10 cells were resisted in *Ythdf2^F/F^*;OT-1 mice but rapidly grown up in the *Ythdf2^CKO^*;OT-1 counterparts (Fig. 2h). To align the initial immune state, an equivalent number of *in vitro*-activated CD8 T cells of *Ythdf2^F/F^*;OT-1 or *Ythdf2^CKO^*;OT-1 origin were transferred into mice inoculated with B16F10-OVA cells. As anticipated, YTHDF2-deficient T cells exhibited inferior antitumor efficacy in this setting (Fig. 2i).

Further, to substantiate the importance of YTHDF2 in ICB-induced T cell immunity, we subjected the above mice to grow MC38 or Hepa1-6 cells, both of which are thought to respond vigorously to anti-PD-1 monotherapy. Nonetheless, unlike *Ythdf2^F/F^* mice, the *Ythdf2^CKO^* littermates produced a compromised response to PD-1 blockade in the MC38 model and gained no aid of killing effect toward the Hepa1-6 tumors (Fig. 2j, k). In keeping with this, anti-PD-1 therapy led to a much lower frequency of tumor-specific or CX3CR1^+^Tim3^+^CD101^-^ T_eff_-like CD8 T cells as well as less cytokine production by these cells in *Ythdf2^CKO^* mice (Fig. 2l and Supplementary Fig. 3f), indicating an indispensable role for YTHDF2 in implementing ICB-elicited T cell functionality.

Together, these observations have indicated that YTHDF2 expression is fundamental for antitumor effector and effector-like CD8 T cells, which constitute natural and ICB-induced immunity, respectively.

### YTHDF2 prevents T cell mitochondrial dysfunction through m^6^A-dependent RNA decay

To understand how YTHDF2 depletion results in T cell dysfunction and to minimize the disturbance posed by the TME, we assessed *in vitro*-activated *Ythdf2^F/F^* and *Ythdf2^CKO^* CD8 T cells under different conditions. Consistent with our *in vivo* results, *Ythdf2*-deficient CD8 T cells yielded a decline in survival and cytokine production as well as a susceptibility to exhaustion, but did not differ in memory T cell differentiation (Supplementary Fig. 4).

In terms of a selective expression pattern, by performing RNA sequencing, we asked whether the abnormalities caused by YTHDF2 loss were rooted in early effector CD8 T cells while doing RNA sequencing. Among the 611 genes upregulated upon YTHDF2 ablation, gene ontology (GO) analysis revealed dominant enrichment for gene sets related to mitochondrial organization and mRNA translation (Supplementary Table 1 and Fig. 3a, b). *Ythdf2^CKO^* CD8 T cells exhibited perturbed mitochondrial membrane potential and accumulated mitochondrial mass and reactive oxygen species (ROS) both *in vitro* and *in vivo* (Fig. 3c–f). We further probed the metabolic phenotype of OVA-activated *Ythdf2^CKO^;OT-1* CD8 T cells. *Ythdf2^CKO^;OT-1* CD8 T cells exhibited decreased extracellular acidification rate (ECAR) and oxygen consumption rate (OCR) (Supplementary Fig. 5a, b) during a mitochondrial stress test. Morphologically, activated *Ythdf2^CKO^;OT-1* CD8 T cells had swollen and fewer mitochondria with disorganized cristae, while *Ythdf2^F/F;^OT-1* CD8 T cells had compact mitochondria with tightly packed cristae (Supplementary Fig. 5c). These observations suggest that YTHDF2 loss-associated redundant mRNA translation and mitochondrial mass resulted in T cell stress, which can explain the susceptibility to exhaustion during chronic TCR stimulation^35^ (Supplementary Fig. 4d). Given the established causal relationship between mitochondrial malfunction and T cell exhaustion^24,36^, YTHDF2 might be critical for ensuring mitochondrial fitness and T cell persistence. Since*Ythdf2^CKO^* CD8 T cells did not preferentially express genes related to programmed cell death, we reasoned that the perturbed cell survival might also be a result of mitochondrial stress (Supplementary Table 1 and Fig. 3b). We then employed N-acetylcysteine (NAC) to scavenge mitochondrial ROS in *in vitro* stimulated CD8 T cells. Nonetheless, no effect on T cell proliferation or cytokine production was observed, but NAC was sufficient to prevent excessive exhaustion and cell death caused by YTHDF2 loss (Supplementary Fig. 5d, e, g and Fig. 3g, h).

**Fig. 3.**
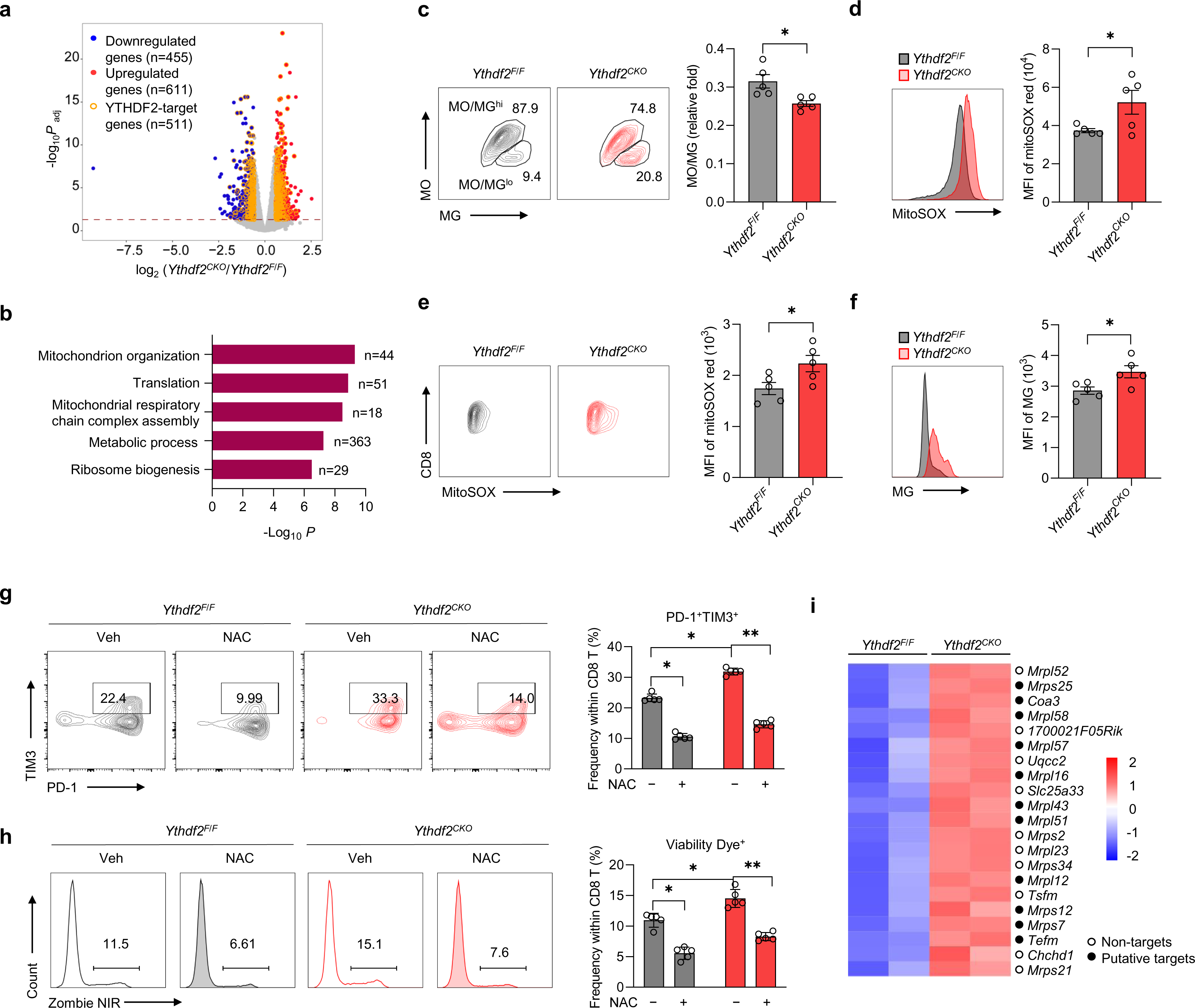
YTHDF2 prevents mitochondrial stress and T cell exhaustion. **a** Volcano plots of genes with differential expression in activated *Ythdf2^F/F^* or *Ythdf2^CKO^* CD8 T cells (anti-CD3/CD28, 5 μg/ml, 24 h) (n = 2 per group), putative YTHDF2 targets that were enriched in both RIP-seq and m^6^A-seq are marked with yellow circles. **b** GO enrichment analysis of upregulated genes in *Ythdf2^CKO^* compared with *Ythdf2^F/F^* CD8 T cells after priming (anti-CD3/CD28, 5 μg/ml, 24 h). **c** Mitochondrial membrane potential and mitochondrial mass was measured by MitoTracker Orange (MO) and MitoTracker Green (MG) staining in activated *Ythdf2^F/F^* and *Ythdf2^CKO^* CD8 T cells (n = 5 per group). Mitochondrial fitness was evaluated according to the MO/MG ratio. **d** Mitochondrial ROS was measured by MitoSOX staining in activated CD8 T cells from *Ythdf2^F/F^* or *Ythdf2^CKO^* mice (n = 5 per group). **e** MitoSOX staining in MC38 tumor-infiltrating CD8 T cells from *Ythdf2^F/F^* and *Ythdf2^CKO^* mice (n = 5 per group). **f** Quantification of the MG MFI in MC38 tumor-infiltrating CD8 T cells from *Ythdf2^F/F^* or *Ythdf2^CKO^* mice (n = 5 per group). **g–h** Quantification of Tim3^+^ PD-1^+^ (g), Zombie NIR^+^ (h) frequencies among primed *Ythdf2^F/F^* and *Ythdf2^CKO^* CD8 T cells (anti-CD3/CD28, 5 μg/ml, 48 h) in the presence of 10 mΜ NAC or veh (n = 5 per group). **i** Heatmap showing the relative expression of representative genes (mitochondrion-related and up-regulated in *Ythdf2^CKO^*) in activated *Ythdf2^F/F^* and *Ythdf2^CKO^* CD8 T cells from RNA-seq data. Putative YTHDF2 targets are depicted by filled circles. Error bars, mean ± s.e.m. \**P* < 0.05; \*\**P* < 0.01. Two-way ANOVA (g, h) or two-tailed unpaired Student’s t-test (c–f).

To elucidate the underlying molecular mechanism, we performed RNA-immunoprecipitation sequencing (RIP-seq) to map the target transcripts bound by YTHDF2 in CD8 T cells. Integrative analyses of RIP-seq, m^6^A-seq and RNA-seq data indicated that 47.9% of the differentially expressed genes caused by YTHDF2 deficiency were eligible for YTHDF2 recognition and m^6^A modification (Fig. 3a and Supplementary Table 2). As potential RNA decay targets, the aforementioned mitochondria-related genes (including mitochondrial ribosomal protein-encoding genes) were frequently found among the YTHDF2- and m^6^A-bound transcripts (Fig. 3i). Noticeably, YTHDF2-RIP-seq and m^6^A-seq identified overlapping peaks on the downstream coding regions of *Coa3*, *Mrpl16*, *Mrps12* and *Tefm* mRNAs (Supplementary Fig. 6a). Consistent with the increased mitochondrial stress, the half-lives of these transcripts were significantly prolonged in the absence of YTHDF2 (Supplementary Fig. 6b).

Similarly, the knockdown of YTHDF2 in Jurkat cells also led to excessive mitochondria-related gene expression and ROS accumulation (Supplementary Fig. 6c–f and Supplementary Table 3). To determine the dependency of this regulatory process on the m^6^A machinery, we exogenously reconstituted YTHDF2 in Jurkat-shYTHDF2 cells. The overexpression of wild-type YTHDF2, but not its inactive mutants that fail in recognition of m^6^A^37^, preserved mitochondrial fitness (Supplementary Fig. 6g–k). However, dampening METTL3, which constructs the m^6^A methylome in T cells^19^, rekindled mitochondrial malfunction despite the presence of abundant YTHDF2 expression (Supplementary Fig. 6l–n). These results demonstrate that the m^6^A machinery is essential for YTHDF2-regulated mitochondrial fitness in T cells.

### YTHDF2 sequesters IKZF1/3 to dictate an active chromatin state in polyfunctional CD8 T cells

Despite identifying the YTHDF2-mediated regulation of CD8 T cell persistence, the mechanism by which YTHDF2 promotes CD8 T cell polyfunctionality remains to be addressed. Interestingly, YTHDF2 depletion was thought to mainly stabilize its target genes, but RNA profiling revealed a comparable number of downregulated genes, which integrally reflected an inactive chromatin state in *Ythdf2^CKO^* CD8 T cells (Supplementary Table 1 and Fig. 4a). Provided that recent studies have uncovered a m^6^A-responsible crosstalk between RNA modification and chromatin regulation^22,23,25^, we tested whether gene transcription was affected by YTHDF2 in early effector CD8 T cells. Consistent with the distinguishing gene signature, compared with *Ythdf2^F/F^* (or *Ythdf2^F/F^*;OT-1) CD8 T cells when shortly activated *in vitro*, *Ythdf2^CKO^* (or *Ythdf2^CKO^*;OT-1) CD8 T cells manifested a remarkable decrease in the abundance of nascent transcripts (Supplementary Fig. 7a, b), hinting that YTHDF2 plays an important role in chromatin remodelling. No genes encoding general epigenetic regulators or transcription factors were found among those putative decay targets (Supplementary Table 2), excluding the possibility of indirect chromatin regulation rooted in the canonical function of YTHDF2. Interestingly, in the presence of an RNA polymerase II-selective inhibitor α-Amanitin^38^, nascent transcripts were significantly reduced in *Ythdf2^F/F^* but not *Ythdf2^CKO^* CD8 T cells (Supplementary Fig. 7c), suggesting that YTHDF2 mainly promotes RNA polymerase II-dependent transcription.

**Fig. 4.**
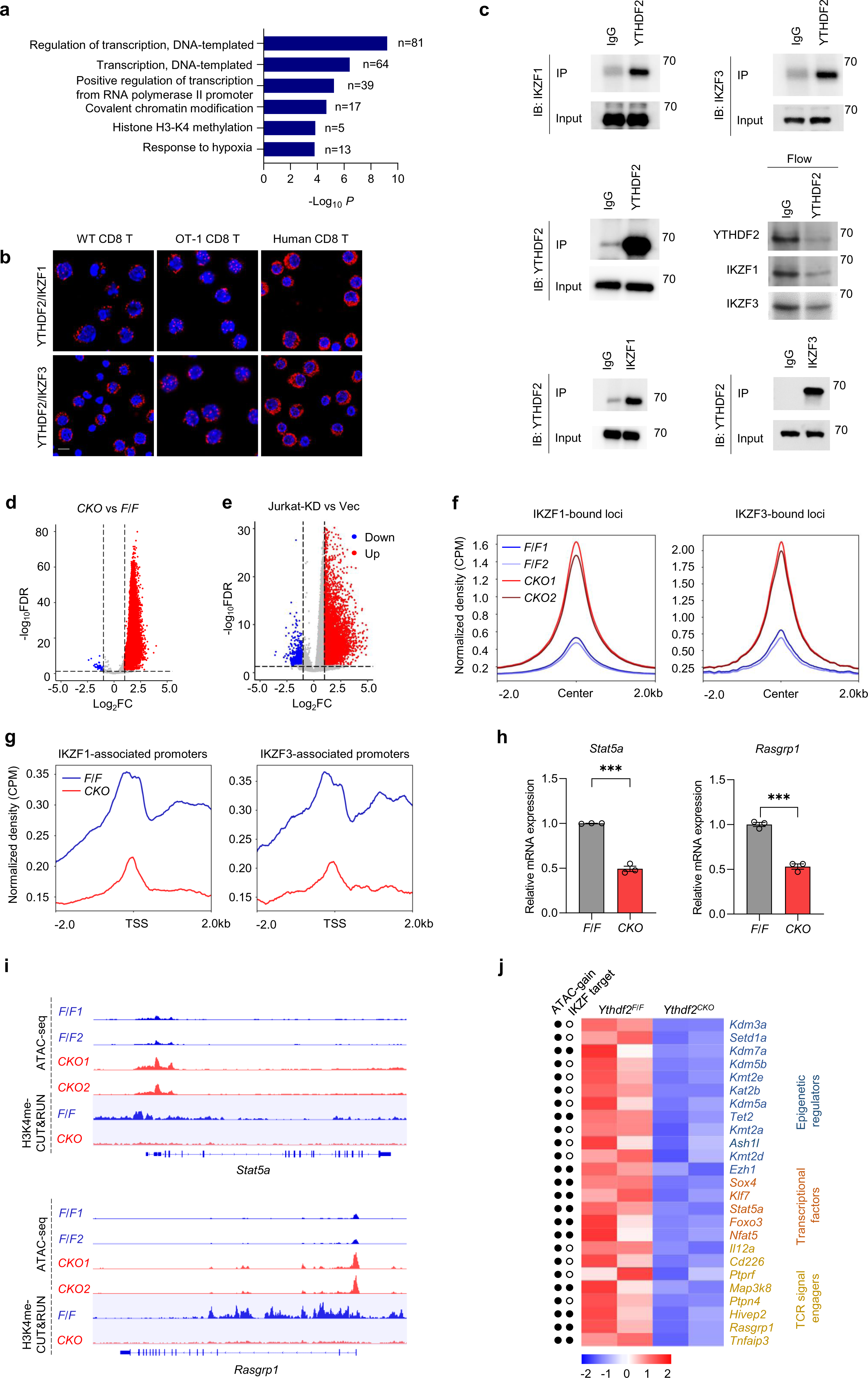
YTHDF2 segregates IKZF1/3 to dictate an active chromatin state in polyfunctional CD8 T cells. **a** GO enrichment analysis of downregulated genes in *Ythdf2^CKO^* compared with *Ythdf2^F/F^* CD8 T cells after priming (anti-CD3/CD28, 5 μg/ml, 24 h). **b** Proximity ligation assay (PLA) analysis of YTHDF2 associated with IKZF1 or IKZF3 in primed mouse (WT or OT-1) and human CD8 T cells stimulated with anti-CD3/CD28 (5 μg/ml, 24 h) or OVA (10 nM, 24 h). Scale bar, 10 μm. **c** Coimmunoprecipitation assays of YTHDF2 associated with IKZF1 or IKZF3 in primed WT CD8 T cells (anti-CD3/CD28, 5 μg/ml, 24 h). **d** Volcano plot of genes with differential chromatin accessibility between activated *Ythdf2^F/F^* and *Ythdf2^CKO^* CD8 T cells (anti-CD3/CD28, 5 μg/ml, 24 h). **e** Volcano plot of genes with differential chromatin accessibility between Jurkat-shCtrl (Vec) and Jurkat-shYTHDF2 (KD) cells. **f–g** ChIP-seq datasets for IKZF1 (GSM1296538) and IKZF3 (GSM803106) in mouse T cells were obtained using Cistrome Data Browser. ATAC-seq (f) or H3K4me CUT&RUN (g) profiles of activated *Ythdf2^F/F^* and *Ythdf2^CKO^* CD8 T cells were represented on IKZF1/3-bound loci or IKZF1/3-associated promoters. **h** *Stat5a* (left) and *Rasgrp1* (right) mRNA levels detected by RT-qPCR in activated *Ythdf2^F/F^* and *Ythdf2^CKO^* CD8 T cells (anti-CD3/CD28, 5 μg/ml, 24 h) (n = 3 per group). **i** ATAC-seq and H3K4me CUT&RUN tracks on the gene loci of *Stat5a* (top) and *Rasgrp1* (bottom) in activated *Ythdf2^F/F^* and *Ythdf2^CKO^* CD8 T cells. **j** Heatmap showing the relative expression of representative genes (down-regulated in *Ythdf2^CKO^)* in activated *Ythdf2^F/F^* and *Ythdf2^CKO^* CD8 T cells from RNA-seq data. Enhanced chromatin accessibility or putative IKZF1/3 binding is depicted by filled circle. Error bars, mean ± s.e.m. \*\*\**P* < 0.001. Two-tailed unpaired Student’s t-test (h).

Nuclear YTHDF2 was known to promote translation initiation of stress-inducible transcripts^27^. However, ribosome profiling indicated no difference in translation efficiency between activated *Ythdf2^F/F^* and *Ythdf2^CKO^* CD8 T cells, even for YTHDF2-targeted and m^6^A-marked transcripts (Supplementary Fig. 7d). To investigate the mechanism underlying YTHDF2-directed transcriptional adaptation, we performed immunoprecipitation followed by mass spectrometry (IP–MS) and identified proteins that were bound to YTHDF2 in the scenario of early T cell activation. Mirroring its potential nuclear functionality, YTHDF2-binding partners included the lymphoid transcription factor Ikaros (IKZF1) and Aiolos (IKZF3)^39^ (Supplementary Table 4). Fitting their roles as transcription repressors, IKZF1 and IKZF3 were expressed at lower levels in activated CD8 T cells than in naïve CD8 T cells (Supplementary Fig. 7e). The results of a proximity ligation assay (PLA) showed that YTHDF2 interacted with IKZF1 and IKZF3 in short-term primed mouse or human CD8 T cells and untreated Jurkat cells (Fig. 4b and Supplementary Fig. 7f). Subsequent coimmunoprecipitation (CO-IP) assays further confirmed these results (Fig. 4c). Of note, upon early CD8 T cell activation, a majority of IKZF1 or IKZF3 protein could be bound to YTHDF2. To probe whether RNA species facilitates the interaction between YTHDF2 and IKZF1/3, endogenous YTHDF2 immunoprecipitants from acutely stimulated CD8 T cells were incubated with RNase or DNase. As shown by Supplementary Fig. 7g, neither RNase nor DNase repressed YTHDF2 binding with IKZF1/3.

To assess whether YTHDF2 affects chromatin openness, we performed an assay for transposase-accessible chromatin with high-throughput sequencing (ATAC-seq). Unexpectedly, YTHDF2 deficiency resulted in enhanced chromatin accessibility in both activated CD8 T cells and Jurkat cells (Fig. 4d, e). Notably, among the ‘ATAC gain’ regions conditioned by YTHDF2 depletion, an analysis of transcription factor (TF) motifs revealed that the ACAGGAAG element, which is capable of binding IKZF1 or IKZF3, was the top hit (Supplementary Fig. 7h, i), which is capable of binding IKZF1 or IKZF3. Using existing chromatin immunoprecipitation sequencing (ChIP-seq) datasets^39,40^, we determined that the specific loci for these two transcription factors selectively displayed greater chromatin accessibility in YTHDF2-deficient T cells than in YTHDF2-competent T cells (Fig. 4f and Supplementary Fig. 7j). Epigenomic mapping through cleavage under targets and release using nuclease (CUT&RUN), however, underscored that the transcription start sites (TSSs) of IKZF1/3-regulated genes in YTHDF2-deficient T cells were marked by much lower amounts of histone H3 lysine 4 methylation (H3K4Me) (Fig. 4g), which is an active form of chromatin modification. These unusual chromatin changes were profoundly observed for genes responsible for TCR signalling, such as *Stat5a*^9^ and *Rasgrp1*^41^, which were confirmed to undergo transcriptional silencing in early activated *Ythdf2^CKO^* T cells (Fig. 4h–j). Genes encoding epigenetic modulators were also widely affected, many of which were simultaneously found to have ‘ATAC gain’ regions predicted for IKZF1/3 binding (Fig. 4j).

Since the protein levels of IKZF1/3 were comparable between *Ythdf2^F/F^* and *Ythdf2^CKO^* CD8 T cells (Supplementary Fig. 8a), we asked whether YTHDF2 deficiency had an impact on IKZF-DNA binding activity. As expected, CUT&RUN assay detected stronger IKZF1/3 binding signals at the promoter regions of genes expressed under a YTHDF2-deficient condition (Supplementary Fig. 8b). As suggested by a recent report^42^, nucleosome occupancy at IKZF motifs could inactivate effector gene transcription without reducing T cell chromatin openness. The molecular basis might be IKZF1/3-mediated recruitment of histone deacetylase (HDAC) complexes^43,44^. Here in an early activation scenario, *Ythdf2^CKO^* but not *Ythdf2^F/F^* CD8 T cells enabled strong IKZF-HDAC1 interaction (Supplementary Fig. 8c), raising the possibility of nucleosome occupancy and chromatin inactivation upon YTHDF2 loss. Together, these data highlight that YTHDF2 loss may result in IKZF1/3-associated transcriptional repression.

### Lenalidomide rescues the polyfunctionality of YTHDF2-deficient CD8 T cells

We next sought to unravel the dependency of YTHDF2 loss-associated T cell malfunction on IKZF1 and IKZF3. As shown in Supplementary Fig. 8d, the knockdown of IKZF1 and IKZF3 restored nascent transcription in Jurkat-shYTHDF2 cells but did not change that in control cells. The myeloma drug lenalidomide has been shown to cause the proteasomal degradation of IKZF1 and IKZF3^45,46^. We asked whether this clinically available drug could restore the effector function of YTHDF2-deficient T cells by targeting IKZF1/3. When lenalidomide was added to short-term activated CD8 T cells or untreated Jurkat cells, it did not obviously enforce T cell function under a YTHDF2-competent condition; in contrast, it restored nascent RNA synthesis as well as the proliferation of *Ythdf2^CKO^* CD8 T cells (Supplementary Fig. 8e–h and Fig. 5a–d), manifesting a context-dependent mode of action. Although there was no effect on preventing T cell exhaustion, lenalidomide profoundly provoked cytokine production by *Ythdf2^CKO^* CD8 T cells (Supplementary Fig. 8i and Fig. 5e). Supporting a retrieved functional state, *Stat5a* and *Rasgrp1* expression in *Ythdf2^CKO^* CD8 T cells returned to normal in the presence of lenalidomide (Supplementary Fig. 8j).

**Fig. 5.**
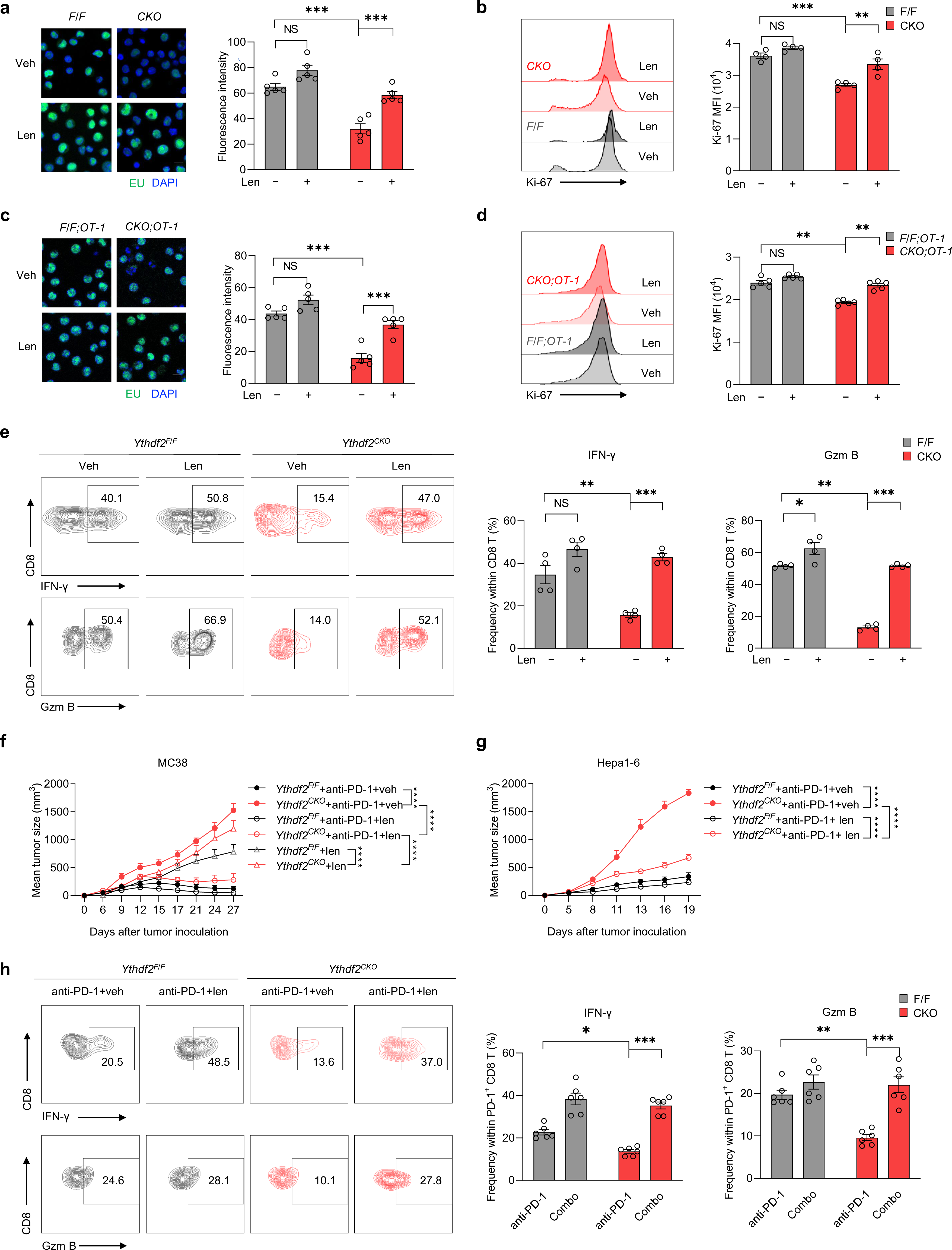
Lenalidomide retrieves the antitumor function of YTHDF2-deficient CD8 T cells. **a** Click-it RNA imaging and analysis of nascent RNA synthesis (green) in *Ythdf2^F/F^* and *Ythdf2^CKO^* CD8 T cells primed (anti-CD3/CD28, 5 μg/ml, 24 h) in the presence of 10 μΜ lenalidomide (len) or vehicle (veh) (n = 5 per group). Scale bar, 10 μm. **b** Quantification of Ki-67 MFI among *Ythdf2^F/F^* and *Ythdf2^CKO^* CD8 T cells primed in the presence of 10 μΜ len or veh (n =4 per group). **c** Click-it RNA imaging and analysis of nascent RNA synthesis (green) in *Ythdf2^F/F^;OT-1* and *Ythdf2^CKO^;OT-1* CD8 T cells primed (OVA, 10 nM, 24 h) in the presence of 10 μΜ len or veh (n = 5 per group). Scale bar, 10 μm. **d** Quantification of Ki-67 MFI among *Ythdf2^F/F^;OT-1* and *Ythdf2^CKO^;OT-1* CD8 T cells primed (OVA, 10 nM, 72 h) (n = 5 per group) in the presence of 10 μΜ len or veh. **e** Quantification of Gzm B, IFN-γ MFI among *Ythdf2^F/F^* and *Ythdf2^CKO^* CD8 T cells primed (anti-CD3/CD28, 5 μg/ml, 48 h) in the presence of 10 μΜ len or veh (n = 4 per group). **f** MC38-bearing *Ythdf2^F/F^* (n = 21) or *Ythdf2^CKO^* (n = 18) mice were treated with anti-PD-1 (250 μg/mouse) and/or len (10 mg/kg) and monitored for tumor growth. **g** Hepa1-6 tumor-bearing *Ythdf2^F/F^* (n = 12) or *Ythdf2^CKO^* (n = 12) mice were treated with anti-PD-1 (250 μg/mouse) and/or len (10 mg/kg) and monitored for tumor growth. **h** Quantification of Gzm B^+^ CD8 T or IFN-γ^+^ CD8 T frequencies within TILs from Hepa1-6-bearing *Ythdf2^F/F^* (n = 6) or *Ythdf2^CKO^* (n = 6) mice treated with anti-PD-1 (250 μg/mouse) and/or len (10 mg/kg) (D13). Error bars, mean ± s.e.m. \**P* < 0.05; \*\**P* < 0.01; \*\*\**P* < 0.001. Two-way ANOVA (a–h).

Taking into consideration of ICB-induced YTHDF2 relocation in effector-like CD8 T cells, we evaluated the dependency of these cells on IKZF1/3 signalling and the therapeutic effect of lenalidomide *in vivo*. Although lenalidomide monotherapy was less efficient than anti-PD-1 therapy when administered to MC38-bearing *Ythdf2^F/F^* mice, it had a slightly better effect on *Ythdf2^CKO^* mice; promisingly, lenalidomide combined with PD-1 blockade largely rescued tumor-eradicating immunity in *Ythdf2^CKO^* mice (Fig. 5f, g). In line with this, flow cytometry analysis showed that combination therapy increased cytokine production by PD-1^+^ *Ythdf2^CKO^* CD8^+^ TILs to a level equivalent to that in the *Ythdf2^F/F^* group receiving anti-PD-1 or combination therapy (Fig. 5h). While it is true lenalidomide is more than an IKZF degrader, these data have supported the likelihood that the unresponsiveness of YTHDF2-deficient T cells was caused by IKZF1/3. Together, we conclude that YTHDF2 may achieve polyfunctionality in effector or effector-like CD8 T cells by preventing IKZF1/3-associated transcriptional repression.

### YTHDF2 relies on m^6^A-bound transcripts for nuclear translocation

Regarding YTHDF2 nuclear relocation in early (re)activated T cells (Fig. 1e–h), we inferred that newly synthesized RNA substrates at this stage might be necessary for YTHDF2 trafficking or acting in the nucleus. In support of this notion, the addition of the transcription inhibitor actinomycin D (ActD) largely blocked the interaction between YTHDF2 and IKZF3 in the T cell nucleus (Fig. 6a). Further, the finding that YTHDF2 was recruited to the m^6^A sites within mRNAs transcribed from its DNA targets raised the possibility of cotranscriptional regulation (Supplementary Fig. 9a and Supplementary Table 5). To determine whether the m^6^A machinery is required for nuclear YTHDF2 distribution and function, we depleted METTL3 in Jurkat cells and mouse T cells. Compared with *Mettl3^F/F^* mice, *Mettl3^CKO^* littermates exhibited accelerated tumor growth, accompanied by decreased proliferation and cytokine production in tumor-infiltrating CD8 T cells (Supplementary Fig. 9b–d). Of importance, YTHDF2 failed to localize to the nucleus of METTL3-deficient CD8 T cells following *in vitro* priming (Fig. 6b). In line with this, METTL3 loss interrupted the binding of YTHDF2 to IKZF3 in both Jurkat cells and activated mouse T cells (Fig. 6c, d). Mutation of a catalytic residue (W386A or W432A) in the hydrophobic pocket of YTHDF2, which specifically recognizes m^6^A, had a similar effect as METTL3 depletion (Fig. 6e and Supplementary Fig. 9e). In parallel, the disruption of METTL3 in YTHDF2-overexpressing cells abrogated its ability to facilitate nascent RNA synthesis and proliferation (Fig. 6f, g). We then explored whether augmented m^6^A mRNA modification could conversely reinforce YTHDF2 nuclear function by repressing the m^6^A demethylase FTO in activated T cells. However, treatment with an FTO inhibitor (FB23-2)^47^ did not promote nascent RNA synthesis regardless of the YTHDF2 concentration (Supplementary Fig. 9f, g), suggesting that increasing m^6^A deposition per se is not sufficient to boost the nuclear function of YTHDF2. These observations suggest that m^6^A recognition might be a prerequisite for YTHDF2 trafficking to the nucleus.

**Fig. 6.**
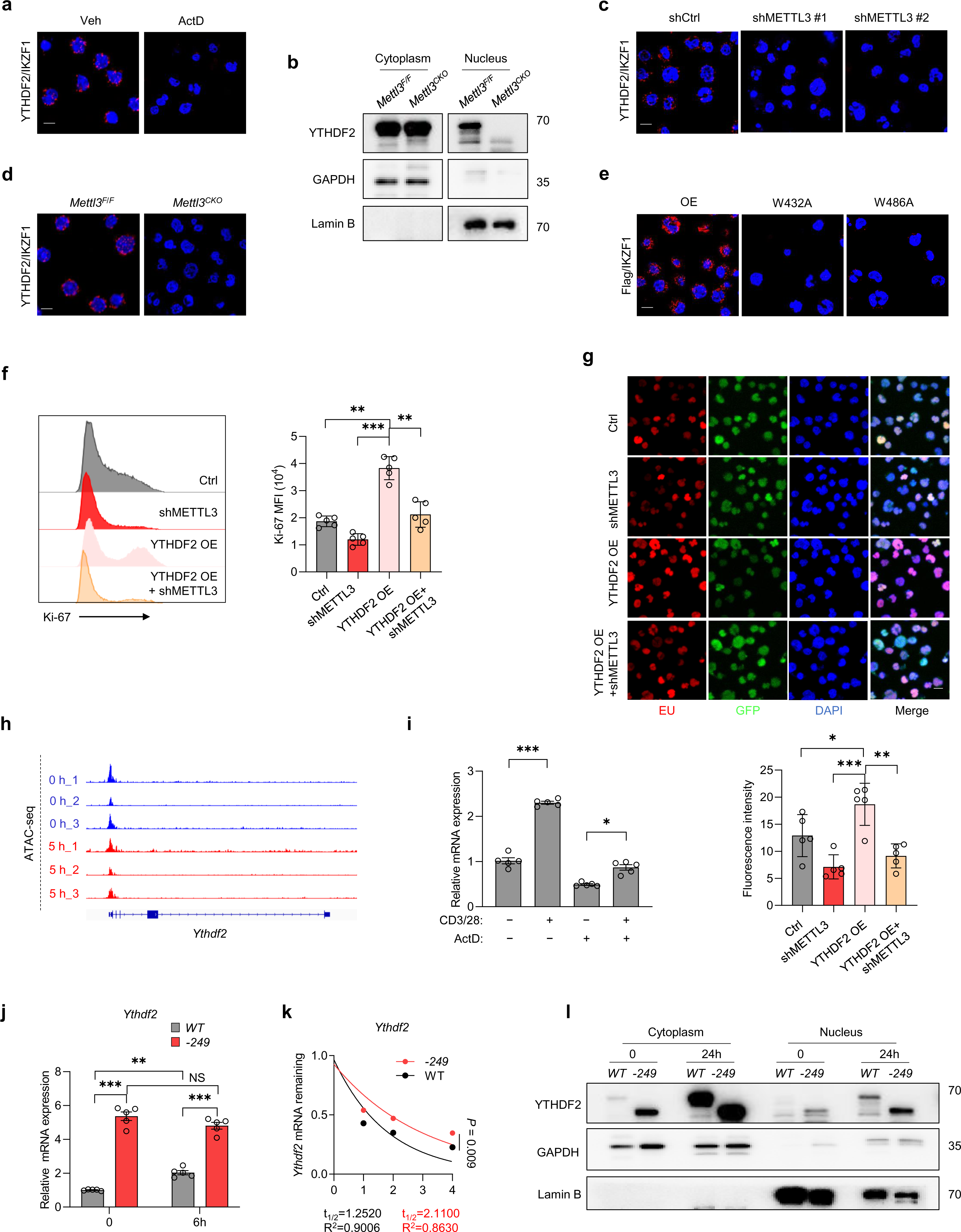
The m^6^A machinery regulates both YTHDF2 relocation and expression. **a** PLA analysis of YTHDF2 associated with IKZF3 in Jurkat cells treated with or without ActD (500ug/ml, 4h). Scale bar, 10 μm. **b** Immunoblotting analysis of YTHDF2 in the cytosol and nucleus of *Mettl3^F/F^* or *Mettl3^CKO^* CD8 T cells stimulated with anti-CD3/CD28 (5 μg/ml, 24 h). **c** PLA analysis of YTHDF2 associated with IKZF3 in Jurkat-shCtrl and Jurkat-shMETTL3 cells. Scale bar, 5 μm. **d** PLA analysis of YTHDF2 associated with IKZF3 in primed *Mettl3^F/F^* or *Mettl3^CKO^* CD8 T cells (5 μg/ml, 24 h). Scale bar, 10 μm. **e** PLA analysis of Flag associated with IKZF3 in Jurkat cells introduced with Flag-tagged WT or mutant YTHDF2. Scale bar, 10 μm. **f** Quantification of Ki-67 MFI among METTL3-knockdown and control Jurkat cells with or without YTHDF2 overexpression (OE) (n = 5 per group). **g** Click-it RNA imaging and analysis of nascent RNA synthesis in METTL3-knockdown and control Jurkat cells with or without YTHDF2 overexpression (OE) (n = 5 per group). Scale bar, 10 μm. **h** ATAC-seq tracks of *Ythdf2* loci on naïve or activated human T cells (anti-CD3/CD28, 5 h) (GSE116696). **i** *Ythdf2* mRNA levels detected by qPCR in CD8 T cells stimulated with or without anti-CD3/CD28 (5 μg/ml) and ActD (500 ug/ml) for 24 h (n = 5 per group). **j** *Ythdf2* mRNA levels detected by qPCR in naïve (0 h) or activated (6 h) WT (n = 5) and *Ythdf2^-249^* (n = 5) CD8 T cells. **K** Naïve *Ythdf2^-249^* and WT CD8 T cells were treated with ActD (500 μg/ml) and RNAs were collected at different time points after ActD treatment. *Ythdf2* mRNA levels were measured using qPCR and represented as mRNA remaining after transcription inhibition (TI) (n = 3 per group). **l** Immunoblotting analysis of YTHDF2 in the cytosol and nucleus of naïve (0 h) or activated (24 h) WT compared with *Ythdf2^-249^* CD8 T cells. Error bars, mean ± s.e.m. NS, no significance; \**P* < 0.05; \*\**P* < 0.01; \*\*\**P* < 0.001. One-way (f, i, g) or two-way ANOVA (j) or non-linear regression (k).

### YTHDF2 expression is posttranscriptionally autoregulated via the m^6^A machinery

Although *Ythdf2* mRNA expression was increased at early timepoints of CD8 T cell priming (Supplementary Fig. 9h), its chromatin accessibility at TSS regions remained unchanged after 5 h of *in vitro* priming (Fig. 6h). The addition of ActD to CD8 T cells did not weaken the fold change in *Ythdf2* mRNA expression, thus excluding a direct transcriptional regulation (Fig. 6i). By incorporating YTHDF2-RIP-seq with m^6^A-seq data from both primed CD8 T cells and untreated Jurkat cells, we noticed that YTHDF2 directly bound to its own mRNA and identified overlapping peaks on its m^6^A-occupied exons (Supplementary Fig. 9i). Therefore, one interpretation of these findings is that YTHDF2 recognizes and destabilizes its cognate mRNA in the cytoplasm; once a portion of YTHDF2 is dissociated and enters the nucleus in response to T cell activation, its mRNA translation can be partially unleashed. To test this hypothesis, we generated a mouse line (*Ythdf2^-249^*) carrying an N-terminal truncated form of YTHDF2, which is unable to translocate to mRNA decay sites but retains the domain for the recognition of methylated RNA^26^. Additionally, this gene engineering strategy did not affect the internal mRNA regions that are able to undergo m^6^A modification. Compared to the wild-type control, naïve *Ythdf2^-249^* CD8 T cells expressed a much higher level of *Ythdf2* mRNA, which did not further increase following 6 h of *in vitro* priming (Fig. 6j). A prolonged *Ythdf2* mRNA lifetime in *Ythdf2^-249^* CD8 T cells further confirmed the preexistence of autoregulated RNA decay (Fig. 6k). *Ythdf2^-249^* CD8 T cells also exhibited increased protein expression in both the cytosol and the nucleus (Fig. 6l). Similarly, at the *Ythdf2* gene locus of *Ythdf2^CKO^* CD8 T cells, RNA-seq captured a higher level of pseudogene expression than that of *Ythdf2* mRNA in *Ythdf2^F/F^* CD8 T cells (Supplementary Fig. 9k). Taken together, these data have shown an unprecedented mode of YTHDF2 autoregulation in CD8 T cells.

### YTHDF2 expression and distribution in human tumor-infiltrating T cells

To explore the clinical relevance of our findings, we reanalyzed single-cell RNA-seq data from human colorectal carcinoma (CRC)^48^ (GSE146771), pancreatic ductal adenocarcinoma (PDA)^49^ (GSE155698), or anti-PD-1-treated hepatocellular carcinoma (HCC)^50^ (GSE206325). We divided the single-cell transcriptomes of CD8^+^ TILs into two groups using a customized polyfunctionality signature score (based on *Ifng*, *Gzma*, *Gzmb*, and *Prf1* gene expression). As shown in Fig. 7a, *Ythdf2* mRNA expression was much greater in CD8 T cells assigned with higher scores. Notably, post-ICB datasets of human melanoma (GSE120575^51^) or HCC (GSE206325) showed that *Ythdf2* level was much greater in CD8^+^ TILs of responders than that of nonresponders (Fig. 7b), suggesting that YTHDF2 is involved in ICB-induced antitumor immunity. Moreover, among the 7 different CD8 T cell subsets from HCC patients who responded to anti-PD-1 therapy, cytotoxic CD8 T cells exhibited the greatest *Ythdf2* expression level (Fig. 7c), which is consistent with the results of our mouse experiments. However, those nonresponder-derived cytotoxic CD8 T cells exhibited a *Ythdf2* level comparable to that of the terminally exhausted population (Fig. 7c).

**Fig. 7.**
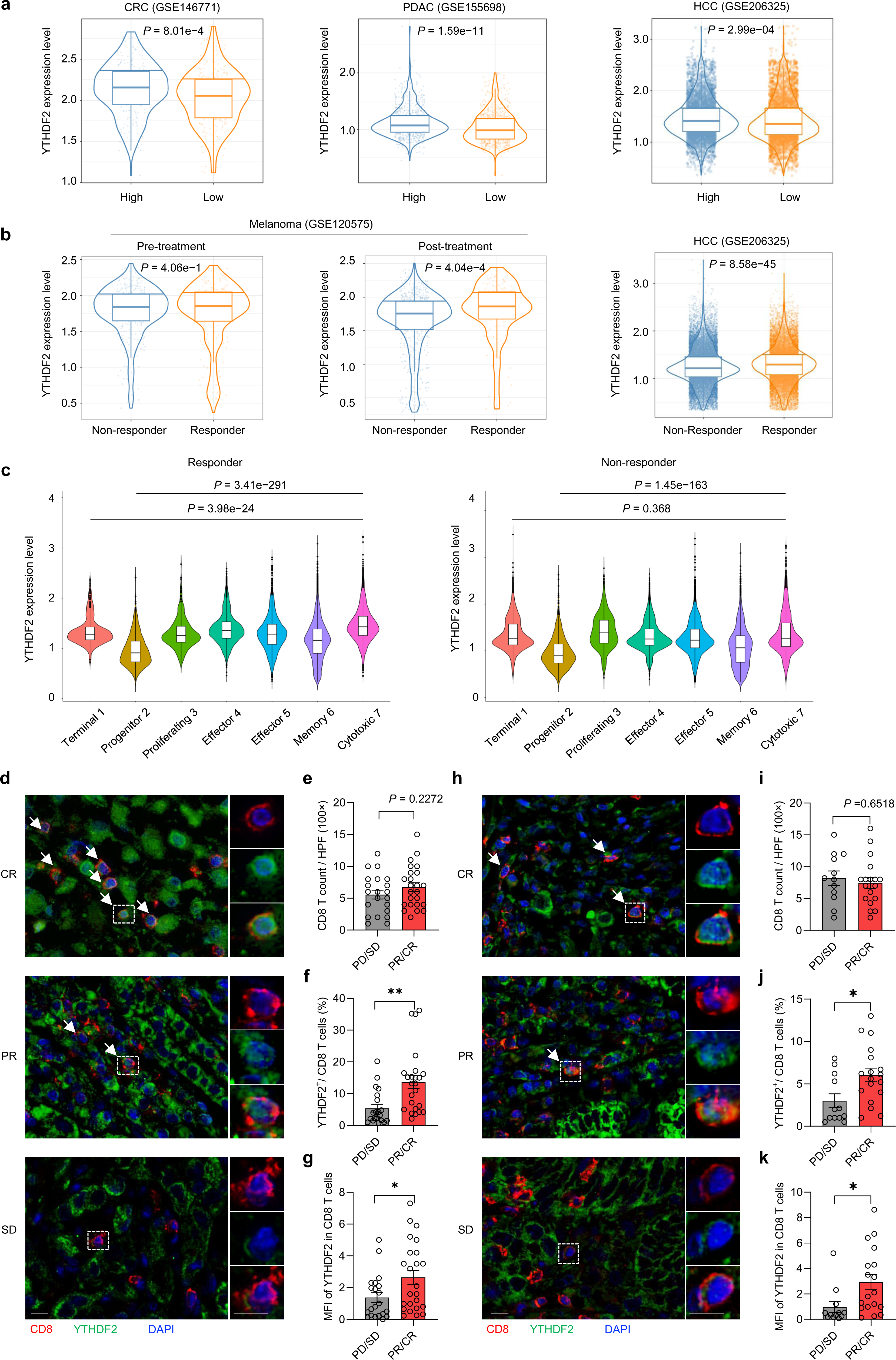
YTHDF2 expression and distribution are associated with T cell function in human cancers. **a** Violin plot comparing*Ythdf2* gene expression levels of CD8 T cells assigned with high or low polyfunctionality signature scores that were derived from single-cell RNA-seq datasets. Left, CRC. Middle, PDAC. Right, HCC. **b** Violin plot comparing pre- or post-treatment *Ythdf2* gene expression levels of CD8 T cells between responders and nonresponders. Left, melanoma. Right, HCC. **c** Violin plot comparing*Ythdf2* gene expression levels among CD8 T cell clusters generated from neoadjuvant anti-PD-1-treated HCC. **d–i** Tissue sections from patients with hepatocellular carcinoma (HCC) (n = 45) or colorectal carcinoma (CRC) (n = 30) were stained for CD8 (red), YTHDF2 (green) and DAPI (blue). **d–h** Representative immunofluorescent images of sections from HCC (d) and CRC (h) patients showing different responses to neo-adjuvant chemo-(immuno-) therapy. A dashed box represents the 4 × enlarged area shown in the bottom panels with separate channels. White arrows point to cells positive for YTHDF2 and CD8. Quantification of CD8 T cells (e, i), frequencies of YTHDF2-positive CD8 T cells (f, j) and quantification of YTHDF2 intensity of CD8 T cells (g, k). CR, complete response; PR, partial response; SD, stable disease; PD, progressed disease. Scale bar, 10 μm. Error bars, mean ± s.e.m. \**P* < 0.05; \*\**P* < 0.01. Two-tailed unpaired Student’s t-test (e–g, i–k).

We further investigated YTHDF2 protein expression using tissue samples from HCC or CRC patients. For those who had not received preoperative treatment, YTHDF2 protein was moderately expressed in cancer cells^32^ but undetectable in CD8^+^ TILs. Results from our clinical trials demonstrated that chemoimmunotherapy significantly improved overall survival (OS) and progression-free survival (PFS) of patients with advanced cancer^52,53^. YTHDF2-expressing CD8 T cells were substantially more common in patients who received neoadjuvant chemoimmunotherapy (Fig. 7d, h), supporting its inducible upregulation concomitant with therapy-induced CD8 T cell infiltration and activation. Although a greater number of CD8^+^ TILs could not distinguish responders from nonresponders to neoadjuvant therapy, YTHDF2 positivity and nuclear accumulation in CD8 T cells were more frequently found in patients who achieved a complete or partial response (Fig. 7d–k). These observations suggest that YTHDF2-expressing CD8 T cells confer therapy-induced antitumor immunity, potentially serving as a clinical indicator of cancer prognosis.

## Discussion

T cell-mediated tumor-eradicating immunity forms the basis of successful cancer treatments^10,20^. Intense efforts have now been invested in elucidating and breaking the T cell-intrinsic barriers to rejuvenation. Differing from effector and memory T cells, exhausted T cells display distinct functional properties, which are attributed greatly to epigenetic and transcriptional mechanisms implicated in T cell differentiation^12,54^. TOX transcriptionally induces exhaustion-associated gene expression and meanwhile recruits chromatin modifiers to repress gene expression involved in T_eff_ differentiation^55^. Similarly, NR4A1 restrains effector gene transcription by shielding AP-1 from its binding chromatin regions and promotes acetylation of histone 3 at lysine 27 (H3K27ac) for activating genes related to T cell dysfunction^56^. Depletion of either TOX or NR4A1 provisions CD8 T cells with an effector phenotype and boosts antitumor immunity. Besides, CD8 T cells acquire DNMT3A-dependnet de novo DNA methylation events upon effector-to-exhaustion transition even when subjected to PD-1 blockade therapy. Co-targeting of this epigenetic program during ICB yields a more effective antitumor response^57^. As characterized in chronic infection and cancer, proliferation-competent T_pex_ cells, which co-express the transcription factor TCF-1 and exhaustion markers, represent the major therapeutic targets of immune interventions^58^, such that epigenetic imprints in this T cell compartment irrevocably affect immunotherapy efficacy. For instance, the SWI/SNF chromatin remodelling complex PBAF^59^ and the NFAT family transcription factor NFAT5^60^ can specifically drive T_pex_ transition to terminal T_ex_ cells, therefore limiting the outcome of T cell-based immunotherapy. In the present work, we embark on investigating RNA epigenetics in terms of both endogenous and ICB-induced T cell immunity, thus uncovering an indispensable role for the m^6^A reader YTHDF2. Aligning with robust antitumor immune responses, YTHDF2 expression is selectively upregulated and redistributed within both terminal T_eff_ and T_eff_-like cells. In accordance, the loss of YTHDF2 in T cells dampens both endogenous and therapy-induced tumor immunity. Unlike previously reported epigenetic events, YTHDF2 depletion does not necessarily affect T cell differentiation, but functionally erodes T cell proliferation and cytokine production. Longitudinal analyses of CD8^+^ TILs suggest that YTHDF2 deficiency can impair effector functionality and durability, thus yielding hyporesponsiveness to anti-PD-1 therapy. Coupled with phenotypic observation and multi-dimensional sequencing, our data further demonstrate that YTHDF2 expression impacts both effector and exhaustion phases through dual mechanisms.

YTHDF2 has been found to exert context-dependent functions in cancer. While verifying the oncogenic roles of m^6^A methylation in some cancer types, several studies postulated the accompanying position of YTHDF2 as an executer for the RNA decay of tumor suppressors^61,62^. In fact, YTHDF2 could also destabilize oncogene-coding mRNAs in a m^6^A-dependent manner^63,64^. As demonstrated in our previous work, YTHDF2 inhibits mouse and human HCC by processing the decay of *Il11* and *Serpine2* mRNAs, which are responsible for inflammation-related cancer progression and metastasis^32^. Otherwise, YTHDF2 overexpressed in leukemic stem cells can decrease the half-life of apoptosis gene *Tnfrsf2*, thereby descending to a cancer-promoting position^65^. With immunobiology appearing as one of the most promising frontiers, the m^6^A machinery in finetuning tumor immunity has been explored^2^. YTHDF2 has been found to dominate immunosuppressive myeloid cell function in both natural and therapy-experienced cancer contexts^8,9^. On the other hand, YTHDF2 promotes NK cell immunity partially by inhibiting the mRNA stability of *Tardbp,* a negative regulator of cell division and proliferation^66^. Uniquely, here we reveal that YTHDF2 dictates both posttranscriptional and transcriptional programs to reinforce the antitumor function of CD8 T cells. In a naïve state, low-level YTHDF2 limits the stability of its cognate encoding mRNA, which harbors bona fide m^6^A sites. Acute activation or rejuvenation signals incite YTHDF2 relocation to the nucleus, albeit temporarily, paving an individual way for its abundant expression in the early phase. In the circumstance of natural or ICB-induced tumor-eradicating immunity, nuclear YTHDF2 combines and curbs the transcriptional repressor IKZF1/3 to safeguard gene transcription and T cell function. Meanwhile, cytoplasmic YTHDF2-mediated mRNA decay can help improve the mitochondrial fitness and persistence of tumor-reactive T cells, opposing their progressive trajectory toward exhaustion. Further, we’ve shown the necessity of m^6^A recognition for YTHDF2 repositioning into the nucleus, which highlights the important crosstalk between RNA modification and chromatin regulation within antitumor CD8 T cells.

RNA-binding proteins have been proven to pervasively participate in transcriptional control^67^. Consistent with this concept, YTHDC1 was found to regulate gene transcription by recruiting a histone modifier or by processing regulatory RNA species in close proximity to active chromatin regions^24,25^. Our data show that YTHDF2 depletion renders T cell chromatin more accessible to IKZF1 and IKZF3 and trapped in an inactive state, which influences the downstream effector genes such as *Stat5a* and *Rasgrp1*. Instead of regulating chromatin modifier transcripts in the cytoplasm, nuclear-localized YTHDF2 directly interacts with repressive transcription factors. Despite this, its relocation depends on m^6^A-labelled nascent RNAs, reminiscent of the previous observation that m^6^A deposition was essential for partitioning the stress-conditioned mRNA-YTHDF2 complexes into phase-separated subcellular compartments^68^. Similar to the perturbed YTHDF2 trafficking in METTL14-deficient mouse embryonic stem cells^68^, nuclear YTHDF2 is hardly detectable when METTL3 is depleted in either activated CD8 T cells or untreated Jurkat cells. Importantly, the genes encoding YTHDF2-bound transcripts largely overlap with the IKZF1/3 targets, suggesting that m^6^A may cotranscriptionally enlist YTHDF2 in chromatin remodelling. As such, m^6^A participates in every step of this non-canonical YTHDF2 signal undertaken in T cells committed to effector or effector-like function.

We previously reported that YTHDF2 expression could be transcriptionally silenced in the hypoxic TME^32^. Here we show its low expression level in human intratumoral T cells might also be explained by the insufficiency of immune response. Only in cancer patients with a better therapeutic response can YTHDF2-expressing CD8 T cells be profoundly detected, which are supposed to address therapy-induced immunity through a positive feedback loop. Otherwise, the paucity of YTHDF2 unmasks an overlooked T cell epigenetic mechanism, posing threat of immunotherapy resistance. Rationally, the immunomodulatory drug lenalidomide, which targets IKZF1/3 for degradation, facilitates ICB-induced rejuvenation of YTHDF2-deficient T cells. Thus, our understanding of the YTHDF2-centered regulatory circuit in antitumor T cells may inspire novel paths for the development of immunotherapies.

## Methods

### Animals

Wild-type (WT) C57BL/6 mice were purchased from Charles River (Beijing, China) for Medical Research. *Ythdf2^flox/flox^* (*Ythdf2^F/F^*) mice in C57BL/6 background were described previously^32^. *Mettl3^flox/flox^* (*Mettl3^F/F^*) mice were kindly provided by Prof. Z. Yin (Jinan University). *dLck^Cre^* and OT-1 TCR transgenic mice were purchased from the Jackson Laboratory. *Ythdf2^F/F^* or *Mettl3^F/F^* mice were then crossed with *dLck^Cre^* transgenic mice to obtain *Ythdf2^CKO^* or *Mettl3^CKO^* mice with *Ythdf2* or *Mettl3* conditionally knocked out in T cells. For animal experiments referring to *Ythdf2^CKO^* or *Mettl3^CKO^* mice, littermate controls with normal YTHDF2 (*Ythdf2^F/F^*) or METTL3 (*Mettl3^F/F^*) expression were used. *Ythdf2^CKO^* mice were also crossed with OT-1 TCR transgenic mice to obtain *Ythdf2^CKO^*;OT-1 mice. *Ythdf2^-2^*^49^ transgenic mice were generated by depleting a 249-amino acid fragment from the N-terminus of *Ythdf2* gene coding region. Correctly targeted mice were determined by PCR and gene sequence. Primers used for genotyping of *Ythdf2^-249^*: Forward–5’-TGTGAATGATGTGGAAGGAA-3’ and Reverse– 5’-CAACAGCAGAGCCTACAA-3’. All mice were maintained under specific pathogen-free conditions. Mice with 8–12 weeks of age were used for all animal experiments. Animals were randomly allocated to experimental groups.

### Cell cultures

Peripheral naïve CD8 T cells were isolated from the mouse spleen by negative selection magnetic beads (STEM CELL). CD8 T cells were cultured in complete RPMI medium (RPMI 1640, 10% FBS, 20 mM HEPES, 1 mM sodium pyruvate, 0.05 mM 2-mercaptoethanol, 2 mM glutamine, 100 μg/ml streptomycin and 100 units/ml penicillin) and stimulated with plate-bound anti-CD3/CD28 in the presence of 10 ng/ml IL-2 (Peprotech) as indicated. To detect T cell proliferation, naïve CD8 T cells were stained with 0.5 μM CellTracker Violet fluorescent dye (Thermo Fisher) in serum-free medium for 20 min at 37 °C, and then washed three times in PBS. Stained cells were activated by plate-bound anti-CD3/CD28 (biolegend) for 24–120 h and detected in the BV421 channel by FACS. To detect T cell activation-induced apoptosis, naïve CD8 T cells were activated by plate-bound anti-CD3/CD28 for different number of hours, then analyzed with an annexin V/propidium iodide kit (BD).

For *in vitro* T cell exhaustion assay^69^, CD8 T cells were seeded at a concentration of 1 million/ml on plates coated with of anti-CD3 (5 μg/ml) and anti-CD28 (2 μg/ml). After 48 h of activation, chronic stimulation was performed using plates coated with anti-CD3 (5 μg/ml). Cells were passaged onto a fresh anti-CD3-coated plate every two days, maintained at 1 million/ml (in the continued presence of 10 ng/mL IL-2), and analyzed via flow cytometry on Day 8.

To induce CD8 T cells with a memory phenotype in vitro, CD8 T cells were activated in a ‘‘transient’’ stimulation condition in which beads were removed after the initial 3-day incubation as reported^28^. CD8 T cells were seeded at a concentration of 1 million/ml in presence of mouse T-activator CD3/CD28 Dynabeads (Thermo Fisher Scientific) and IL-2 (10 ng/ml) at 1:1 beads-to-cells ratio. After 2 days, cells were split 1:2. Beads were removed on day 3 and cells were maintained in culture for 6 days (with fresh media added every 2 days) in the presence of 10 ng/ml IL-2. Cells were analyzed via flow cytometry on Day 9.

To obtain mouse tumor-infiltrating CD8 T_pex_, B16-OVA tumor-derived single cell suspensions were stained with Zombie NIR and then sorted for live CD45^+^CD8^+^PD1^+^Tim3^-^SLAMF6^+^ cells by the BD FACS Aria II Cell Sorter. Sorted T_pex_ were cultured in U-bottom plates and stimulation assays were performed using T-activator CD3/CD28 Dynabeads and IL-2 (10 ng/ml) at 1:2 beads-to-cells ratio with 10 μg/ml anti-PD-1 (RMP1-14, BioXCell) or cIg for 48 h.

Human peripheral blood mononuclear cells (PBMCs) were isolated from three healthy volunteers by gradient centrifugation with Lymphoprep (STEM CELL). CD8 T cells were then purified by EasySep Human CD8 T Cell Isolation Kit (STEM CELL). For *in vitro* T cell activation experiments, cells were plated at 1 million/ml in the presence of Human T-Activator CD3/CD28 Dynabeads (Thermo Fisher Scientific) at 1:1 beads-to-cells ratio supplemented with 30U/ml human IL-2 (Peprotech).

The MC38 (mouse colon adenocarcinoma) cell line was originally from Prof. Y.-X. Fu laboratory (University of Texas Southwestern Medical Center). The B16F10 (mouse melanoma) cell line (ATCC, CRL-6475) was purchased from the American Type Culture Collection and the B16F10-OVA cell line was generated by EGFP-OVA (SIINFEKEL) lentivirus transduction. Mouse hepa1-6 (hepatoma cells) and human Jurkat (Clone E6-1, T lymphoblast) cells were purchased from the Cell bank of the Chinese Academy of Sciences.

MC38, Hepa1-6, B16F10, B16F10-OVA, Jurkat cells were grown in Dulbecco’s modified Eagle’s medium (DMEM) (Invitrogen) or RPMI-1640 medium (Invitrogen) supplemented with 10% fetal bovine serum (FBS) (Gibco), 10 mM HEPES (Gibco) and 1% Penicillin/Streptomycin (Gibco). All cell lines were maintained at 37°C, 5% CO_2_ and routinely tested negative for Mycoplasma.

Short hairpin RNAs (shRNAs) targeting human *Mettl3*, *Ikzf1* and *Ikzf3* were used to generate gene knockdown in Jurkat cells. Target sequences are listed in Supplementary Table 6. Relevant shRNA-expressing lentiviruses were produced by Obio technology or Gene Chem. YTHDF2-overexpressing or -mutant (the m^6^A recognition sites W432 and W486 were mutated into A) lentiviruses were designed and synthesized as previously described^32^. Briefly, Jurkat cells were seeded 1 × 10^6^/ml in a 12 well-plate. HitransG P transfection reagent (GENE) and the corresponding lentiviruses were added. After centrifugation at 1,200 rpm, 37°C for 60 min, cells were incubated at 37°C, 5% CO_2_ overnight then culture medium was changed. To obtain stably transfected clones, these cells were treated with puromycin (3 μg/ml) for 1 week and maintained in 1 μg/ml. The knockdown or overexpression efficiency was confirmed by quantitative PCR and western blot analysis before the cells were used for subsequent experiments.

In some settings, Jurkat cells were treated with 100 μΜ lenalidomide (Selleck) or vehicle for 24 h. Primed CD8 T cells were treated with 10 μΜ lenalidomide or vehicle for 24, 48 or 72 h as indicated. For FTO inhibition, Jurkat cells or primed CD8 T cells were treated with10 μΜ FB23-2 (Selleck) or vehicle for 72 h. To neutralize ROS, primed CD8 T cells were treated with 10 mΜ NAC(Sigma) or vehicle for 48 or 72 h as indicated. To selectively inhibit Pol II, primed CD8 T cells were treated with 2 μg/mL α-amanitin (MCE) or vehicle for 12 h as indicated.

### Tumor growth and treatments

MC38 (1 × 10^6^), Hepa1-6 (1 × 10^6^), B16F10 (5 × 10^5^) or B16-OVA (1 × 10^6^) tumor cells were injected subcutaneously (s.c.) into the right flank of mice. Tumor growth was monitored every 2 or 3 days. Tumor volumes were measured by length (a) and width (b) and calculated as tumor volume = ab^2^/2.

For anti-PD-1 treatment, MC38 or Hepa1-6 tumors were allowed to grow for five or six days then intraperitoneally (i.p.) injected with 250 μg/dose anti-PD-1 (RMP1-14, BioXCell) or control IgG (cIg). Anti-PD-1 or cIg was given on days 6, 9,12, 15 for MC38 tumors while on days 5, 8,11, 14 for hepa1-6 tumors. For adoptive cell transfer therapy, B16F10-OVA (1 × 10^6^) tumor cells were s.c. injected into the right flank of C57BL/6 WT mice (female, 8 weeks). On day 6, tumor-bearing mice were randomly divided into three groups (n = 5) and intravenously injected with either PBS or 1 × 10^6^ OVA-primed (72 h) OT- 1 CD8 T cells from *Ythdf2^F/F^;OT-1* or *Ythdf2^CKO^;OT-1*. Tumor growth was monitored every 2 or 3 days from day 6. For in vivo lenalidomide treatment, tumor-bearing mice were i.p. injected once daily with 10 mg/kg lenalidomide^70^ (Selleck) dissolved in DMSO and diluted in 100 μl PBS or with DMSO in 100 μl PBS.

For antibody-mediated T cell depletion, 200 µg anti-CD4 (BioXCell, BE0003-1, clone GK1.5) or anti-CD8 (BioXCell, BE0061, clone 2.43) was given by i.p. 3 days before MC38 tumor inoculation and on day 1, 2, 4, 8, 12, 16 and 19, relative to tumor injection (day 0).

### Patient samples

Human HCC tissue specimens were collected from 45 patients receiving surgery with informed consent at Sun Yat-sen University Cancer Center (SYSUCC) from 2019 to 2021, who had been administrated with neo-adjuvant chemo-(immuno-) therapy (regional chemotherapy using a FOLFOX (oxaliplatin, leucovorin, and fluorouracil) regimen, supplemented with or without anti-PD-1 therapy). Human CRC tissue specimens were collected from 30 patients receiving surgery at SYSUCC from 2016 to 2019, who had been administrated with neo-adjuvant chemotherapy (systemic chemotherapy using a FOLFIRI (irinotecan, leucovorin, and fluorouracil) regimen). All patients were followed up on a regular basis. The study was approved by the Medical Ethics Committee of SYSUCC. Written informed consent was obtained from the patients who provided samples. All patients had a histological diagnosis of HCC or CRC.

### Flow cytometry

Tumors, tumor draining lymph nodes, livers, peripheral blood and spleens were harvested from mice as indicated in figure legends. Tumors were sliced into small pieces and put into a gentleMACS C Tube (Miltenyi) containing 100 ml Enzyme D, 50 ml Enzyme R, 12.5 ml Enzyme A (Miltenyi) and 2.35 ml RPMI 1640. The C tube was then processed on a gentleMACS Octo Dissociator with Heaters (Miltenyi) for 30 min. The resulting cell suspension was passed through a 70-mm cells strainer (Miltenyi), then washed with PBS buffer containing 0.04% BSA. Single cell suspensions of spleens were depleted of erythrocytes. Cells were re-suspended in staining buffer (PBS with 2% FBS and 1 mM EDTA). To block mouse Fc receptors, cells were incubated with anti-CD16/CD32 antibody (BD) for 10 min. Subsequently, specific antibodies for cell surface epitope staining were added and staining was continued for 30 min at 4°C in the dark. For mitochondrial staining, cells were incubated with 25 nM MitoTracker Orange (ThermoFisher), 50nM MitoTracker Green (ThermoFisher) and 5 uM MitoSOX red (ThermoFisher) in RPMI with 2% FBS for 30 min at 37 °C after staining surface markers. For intracellular staining, cells were stimulated ex vivo with Cell Stimulation Cocktail plus protein transport inhibitors (eBioscience) for 4 h before surface staining^71^. Following incubation, cells were washed twice with buffer before proceeding to intracellular staining. Cells were then fixed and permeabilized using the Foxp3/ transcription Factor Staining Buffer Set (eBioscience) according to the manufacturer’s protocol and stained with intracellular antibodies or respective isotype antibodies. Cells were analyzed by the Cytek Aurora (Cytek) or CytoFLEX (Beckman) machine. Analysis of flow cytometry data was performed using Flowjo 10.7.1 (Treestar). Dead cells stained by live dead blue (eBioscience) or Zombie Aqua (BioLegend) were excluded from analysis. Gating was confirmed with fluorescence-minus-one (FMO) controls for low-density antigens.

### RIP-seq

RIP-seq was conducted in accordance with a previously reported protocol with minor modifications^26^. Mouse activated CD8 T cells (anti-CD3/CD28, 5 μg/ml, 24 h) or Jurkat-YTHDF2 OE cells were collected then the pellet was treated with cell lysis buffer. The 10% lysis sample was saved as input, 80% was used in immunoprecipitation reactions with anti-YTHDF2 (Abcam) or anti-Flag (CST) antibody, and 10% was incubated with rabbit IgG (CST) as a negative control. The RIP step was performed by using Epi RNA immunoprecipitation kit (Epibiotek) following the manufacturer’s protocols. RNA was then extracted using TRIzol reagent (Invitrogen). Input and immunoprecipitated RNA of each sample were used to generate the library using a TruSeq stranded mRNA sample preparation kit (Illumina). Libraries quality were determined on Qseq100 Bio-Fragment Analyzer (Bioptic). The strand-specific libraries were sequenced on Illumina Novaseq 6000 system with paired-end 2 x 150 bp read length.

### m^6^A-seq

Total RNA in CD8 T cells or Jurkat cells was extracted by using TRIzol Reagent. DNase I (Invitrogen) treatment was adopted to remove DNA contamination. Additional phenol-chloroform isolation and ethanol precipitation treatments were performed to remove enzyme contamination. Following meRIP-Seq was carried out as previously described^72^. Briefly, 20 μg purified RNA was fragmented into ∼200 nucleotide-long fragments by incubating in magnesium RNA fragmentation buffer for 6 min at 70°C. The fragmentation was stopped by adding EDTA. Then, RNA clean and concentrator-5 kit (Zymo) was used to purify fragmented total RNA. Next, m^6^A immunoprecipitation was performed by using Epi m^6^A immunoprecipitation kit (Epibiotek). Fragmented total RNA (Input) and immunoprecipitated RNA (IP) were subjected to library construction by using Epi mini longRNA-seq kit (Epibiotek) according to the manufacturer’s protocols. Briefly, reverse transcription was performed using random primers and the ribosome cDNA was removed after cDNA synthesis using probes specific to mammalian rRNA. The directionality of the template-switching reaction not only preserves the 5’ end sequence information of RNA but the strand orientation of the original RNA. Libraries for immunoprecipitated RNA were PCR amplified for 18 cycles. The quality of libraries was determined on Qseq100 Bio-Fragment Analyzer (Bioptic). The strand-specific libraries were sequenced on Illumina Novaseq 6000 system with paired-end 2 x 150 bp read length.

### LC–MS/MS quantification of m^6^A

Total RNA of naïve or activated CD8 T cells (anti-CD3/CD28, 5 μg/ml, 24 h) was extracted by using TRIzol Reagent. Quantification of m^6^A in mRNAs was carried out as previously described^32^.100 ng of mRNA was digested by nuclease P1 (NEB) in 25 μl of buffer containing 25 mM NaCl, and 2.5 mM ZnCl_2_ at 42 °C for 2 h, followed by the addition of NH_4_HCO_3_ and alkaline phosphatase and incubation at 37 °C for 2 h. The sample was then filtered (0.22 μm, Millipore) and injected into the LC-MS/MS. The nucleosides were separated by reverse-phase ultraperformance liquid chromatography on a C18 column using an Agilent 6410 QQQ triple-quadrupole LC mass spectrometer in positive electrospray ionization mode. The nucleosides were quantified by using the nucleoside-to-base ion mass transitions of 282 to 150 (m^6^A) and 268 to 136 (A). Quantification was carried out by comparison with a standard curve obtained from pure nucleoside standards run with the same batch of samples. The m^6^A/A ratio was calculated based on the calibrated concentrations.

### Bulk RNA-seq

RNA-seq was performed as previously described^32^. Briefly, RNA was isolated from CD8 T cells or Jurkat cells using TRIzol for subsequent RNA library construction. The libraries were sequenced on Illumina nova 6000 in a 150-bp pair-end run (PE150).

### ATAC-seq

ATAC libraries were generated as described with minor modifications^73^. In brief, mouse CD8 T cells or Jurkat cells were harvested and counted. Nuclei from 50,000 cells were isolated using a lysis solution composed of 10 mM Tris-HCl, 10 mM NaCl, 3 mM MgCl_2_, and 0.1% IGEPAL CA-630. Immediately after cell lysis, nuclei were pelleted in low-bind 1.5-ml tubes and resuspended in transposition mix (10 µl 5 x TD buffer, 5 µl Tn5 transposase, 35 µl nuclease-free water). The transposition reaction was performed at 37°C for 45 min. DNA fragments were purified from enzyme solution using Zymo DNA Clean and Concentrator TM -5 kit (Zymo). Libraries were barcoded (Nextera Index Kit, Illumina) and amplified with NEBNext High Fidelity PCR Mix (New England Biolabs). Size selection of the PCR product were performed by using DNA clean beads (Epibiotek). The quality of libraries was determined on Qseq100 Bio-Fragment Analyzer (Bioptic) and sequenced on Illumina Novaseq 6000 system with paired-end 2 x 150 bp read length.

### Ribo-seq

Ribosome profiling libraries were prepared as described with minor changes^74^. Briefly, CD8 T cells were exposed to cycloheximide (CHX, 100 μg/ml) for 15 min, washed twice with 5 ml cold PBS with CHX (100 μg/ml), pelleted, and lysed in Lysis Buffer (20 mM Tris-HCl, pH 7.8, 100 mM KCl, 10 mM MgCl_2_, 1% Triton X-100, 2 mM DTT, 100 μg/ml cycloheximide, 1:100 protease inhibitor, 40 U/ml SUPERasin). The following Ribo-Seq experiment were performed by using Epi^TM^ Ribosome Profiling Kit (Epibiotek) according to the manufacturer’s protocols. Released RNA fragments were purified using Zymo RNA Clean&Concentrator^TM^-5 kit (Zymo) and ribosomal RNA was deleted by using RiboRNA Depletion Kit (Epibiotek). Ribosome protected fragments (RPF) were recovered by using Zymo RNA Clean&Concentrator^TM^-5 kit (Zymo). RNA fragments were prepared into libraries using a QIAseq miRNA Library kit (Qiagen). Size selection of the library products were performed by using Native-PAGE electrophoresis to capture fragments range from 178-180bp. Libraries quality were determined on Qseq100 Bio-Fragment Analyzer (Bioptic) and sequenced on Illumina Novaseq 6000 system with single-end 1 x 75 bp read length.

### Immunoprecipitation (IP)

Cells were washed with 2 ml of phosphate-buffered saline twice and then lysed with IP lysis buffer (Beyotime). After incubation on ice for 15 min and centrifugation at 12,000 rpm at 4°C for 20 min, the supernatant was saved and the protein concentration was determined with the BCA assay (ThermoFisher). Proteins were either directly analyzed by immunoblotting as input or used for immunoprecipitation analysis. Briefly, the proteins were first incubated with the corresponding antibodies (anti-YTHDF2 (Abcam, ab246514), anti-Ikaros (CST, 14859) or anti-Aiolos (CST, 15103)) overnight and then mixed with Protein G beads (MCE) and incubated for 2 more hours. The beads were collected with a magnetic stand (ThermoFisher) and washed five times with Wash Buffer. After the final wash, the beads were resuspended and heated in loading buffer and the supernatant was electrophoresed through SDS-PAGE.

### Mass spectrometry (MS)

Protein lysates from naïve and activated CD8 T cells were immunoprecipitated with anti-YTHDF2 (Abcam, ab246514) antibody, separated by SDS-PAGE, and finally visualized with Coomassie brilliant blue staining. Protein in-gel digestion and nano-HPLC MS/MS were carried out as described^75^.

### CUT&RUN

CUT&RUN was carried out as previously described^76^. Briefly,10^5^ primed *Ythdf2^F/F^* and *Ythdf2^CKO^* CD8 T cells (anti-CD3/CD28, 5 μg/ml, 24 h) were washed and bound to concanavalin A-coated magnetic beads, then permeabilized with Wash Buffer (20 mM HEPES pH 7.5, 150 mM NaCl, 0.5 mM spermidine and protease inhibitor cocktails from Sigma-Aldrich) containing 0.05% digitonin (Dig Wash), and incubated with anti-H3K4Me (Active Motif, 39635), anti-Ikaros (CST, 14859) or anti-Aiolos (CST, 15103) overnight at 4°C. Cell-bead slurry was washed twice with 1 ml Dig Wash, incubated with Protein A-MNase (pA-MN) for 1h at 4C, then washed twice more with Dig Wash to remove unbound PA/G-MNase protein. Slurry was then placed on a pre-cooled metal block and incubated with cold Dig Wash containing 2 mM CaCl_2_ to activate pA-MN digestion. After 30 min incubation, one volume of 2 x Stop Buffer (340 mM NaCl, 20 mM EDTA, 4 mM EGTA, 0.02% Digitonin, 50 μg/ml glycogen, 50 μg/ml RNase A, 4 pg/ml heterologous spike-in DNA) was added to stop the reaction, and fragments were released by incubating the tubes on a heat block at 37C for 30 min. Samples were centrifuged 5 minutes at 16000xg at 4°C, and supernatant was recovered and DNA extracted via phenol-chloroform extraction and ethanol precipitation. Extract DNA was processed for library generation using the QIAseq Ultralow Input Library Kit (QIAGEN) following the manufacturer’s protocol. Libraries quality were determined on Qseq100 Bio-Fragment Analyzer (Bioptic) and sequenced on Illumina Novaseq 6000 system with paired-end 2×150 bp read length.

### Proximity Ligation Assays (PLA) Fluorescence Assay

The DuoLink In Situ Red Starter Kit Mouse/Rabbit (Sigma-Aldrich) was used to detect interacting proteins. The assay was performed according to the manufacturer’s instructions. Glass bottom cell culture dishes were treated with poly-lysine (Sigma-Aldrich) at 37°C for 4 hours. Then cells were seeded to the culture dishes and settled for 15 min. Cells were fixed with 4% paraformaldehyde solution for 20 min. Then the dishes were permeabilized with 0.05% Triton X-100 and blocked with Duolink Blocking Solution in a pre-heated humidified chamber for 60 min at 37°C. The primary antibodies (anti-YTHDF2 (Abcam, ab246514), anti-DYKDDDDK Tag antibody (Cell Signaling Technology, 14793), anti-Ikaros (Proteintech, 66966), anti-Aiolos (Lesding Biology, AMM16470VCF), and anti-HDAC1 (Proteintech, 66085)) were added to the dishes and incubated overnight at 4°C. Then the dishes were washed with Wash Buffer A and subsequently incubated with the PLA probes for 60 min, the Ligation-Ligase solution for 30 min, and the Amplification-Polymerase solution for 100 min in a pre-heated humidified chamber at 37°C. Before imaging, the dishes were mounted with a cover slip using Duolink In Situ Mounting Medium with DAPI. Fluorescence images were acquired using a ZEISS LSM880 with fast airyscan confocal microscope.

### Nuclear and cytoplasmic protein extraction

Mouse CD8 T cells or Jurkat cells were harvested and washed twice in PBS. Cytoplasmic and nuclear fractions were separated using the Minute Cytoplasmic and Nuclear Fractionation kit (Invent) according to the manufacturer’s instructions. Briefly, cells were lysed in the cytoplasmic lysis buffer on ice for 5 min and then centrifuged at 12000 rpm for 5 min at 4°C. The supernatant containing the cytoplasmic proteins were harvested. The pellet containing the cell nucleus were wash with ice-cold PBS for 5 times. Then the pellet was lysed with the nuclear lysis buffer on ice for 40 min and with violent vortex for 15s every 10 min. The nuclear lysate was centrifuged at 12,000 rpm for 5 min and the supernatant containing the nuclear proteins were harvested. The concentrations of both fractions were normalized with BCA assay and subjected to western blot analysis.

### Western blot analysis

Cells were lysed on ice for 15 min using lysis buffer (Beyotime) supplemented with a protease inhibitor cocktail (ThermoFisher). The cell lysate was centrifuged at 12,000 rpm at 4°C for 20 min. The protein concentrations were normalized with a BCA assay kit (ThermoFisher). Equivalent proteins were loaded into 10% SDS-PAGE Gel and transferred to PVDF membranes (Life Technologies). Membranes were blocked for 1 h in TBST buffer with 5% skim milk and then incubated with primary antibodies in the blocking buffer at 4°C overnight. After being washed three times in TBST, membranes were incubated with secondary antibodies for 1 h at room temperature. The quantitative densitometry of immunoblots was analyzed by using Image J software. Relevant antibodies are listed in Supplementary Table 6.

### Nascent RNA labeling assay

CD8 T cells or Jurkat were cultured on pre-coated glass over slides. A nascent RNA synthesis assay was conducted using Click-It RNA Imaging Kits (Invitrogen) following the manufacturer’s protocols. Images were captured with LSM 880 (Zeiss) confocal and intensity of signal was quantified using Zen 2.6 software (Zeiss).

### Quantitative PCR (qPCR)

RNAs were extracted from primary CD8 T cells and Jurkat cells by using RNA-Quick Purification Kit (ESscience), according to the manufacturer’ protocol, and were reverse transcribed using the PrimeScript RT Master Mix (Takara). The GoTaq qPCR Master Mix (Promega) was used to perform quantitative real-time PCR on the LightCycler 480 System (Roche). The primer sequences are listed in Supplementary Table 6.

### Measurement of RNA lifetime

Cells were treated with Act D (500 μg/ml, MCE) for 1, 2, or 4 h. Untreated cells were used as 0 h. Cells were collected at the indicated timepoints. The total RNA was purified by EasySep™ Total Nucleic Acid Extraction Kit (STEM CELL) with an additional DNase-I digestion step (Invitrogen). The quality of the total RNA was assessed using a Bioanalyzer 2100 instrument and the RNA 6000 Nano Assay Kit (Agilent). RNA quantities were determined using qPCR.

### Immunofluorescent Staining

YTHDF2 and CD8 in mouse samples or human specimen were detected by Tyramide SuperBoost kits (Alexa Fluor 488-labeled tyramide, Cy3-labeled tyramide) (ThermoFisher), according to the manufacturers’ protocol. Briefly, paraffin-embedded tissue specimen was first dewaxed at 70°C for 20 min. After antigen retrieval and blocking, tissue sections were incubated with corresponding primary antibodies overnight at 4°C and then incubated with poly-HRP-conjugated secondary antibody and Alexa Fluor tyramide reagent. Finally, HRP reaction was stopped and the tissue sections were multiplexed for second and third signal detection. Nucleus was counterstained with DAPI. Whole slide overview images at 40x magnification were obtained using Pannoramic MIDI (3DHISTECH).

### Seahorse metabolic assay

Seahorse assay was performed to measure OCR and ECAR of primed *Ythdf2^F/F;^OT-1* or *Ythdf2^CKO^;OT-1* CD8 T cells. CD8 T cells were washed in assay media (XF RPMI medium pH 7.4 (Agilent)) and seeded in a 96-well Seahorse Cells Culture Plate (Agilent) in a non-CO2 incubator at 37 °C for 40 min. OCR and ECAR were measured by a Seahorse XFe96 Extracellular Flux Analyzer (Agilent) following the manufacturer’s instructions. During a mito-stress assay, cells were treated with oligomycin (1.5μM, Sigma-Aldrich), carbonylcyanide-4-(trifluoromethoxy) phenylhydrazone (FCCP, 1.5 μM, Sigma-Aldrich), rotenone (0.5 μM, Sigma-Aldrich) and antimycin A (0.5 μM, Sigma-Aldrich). During a glycolysis assay, cells were treated with glucose (10 mM, Sigma-Aldrich), oligomycin (1 μM, Sigma-Aldrich) and 2-DG (50 mM, Sigma-Aldrich). Each condition was performed with 3–6 replicates in a single experiment. OXPHOS and glycolysis were calculated according to the previous report^77^.

### Transmission electron microscopy

Cell pellets were fixed in 2.5% glutaraldehyde for 4h at 22°C. Following pre-fixation, samples were washed in PBS and post fixed in 1% osmium tetroxide for 1h at 22°C. After several washes in PBS and dehydration in acetone, samples were embedded in Epon. Ultrathin sections of 100 nm were prepared on a Leica EM UC7 Ultramicrotome (Leica Microsystems) and stained with uranyl acetate and lead citrate. Images of mitochondria morphology were captured using a Tecnai G2 Spirit transmission electron microscope (FEI Company).

### Processing of single-cell RNA sequencing data

Human single-cell RNA sequencing datasets (GSE206325^50^, GSE146771^48^, GSE155698^49^, GSE120575^51^) were pretreated by R (v4.2.2). The metadata was loaded and pre-processed using the R package Matrix (v1.4-1). For the analysis of tumor-infiltrating CD8 T cells in the violin plot, we excluded cells that met the criteria as reported^78^. Single-cell data processing was carried out using the R package Seurat (version 4.3.0). The defining gene sets for the polyfunctionality score of CD8 T cells include *Ifng*, *Gzma*, *Gzmb*, and *Prf1*. The scRNA-seq gene set functional score was generated using the R package Seurat (version 4.3.0) function ‘AddModuleScore’^79–82^. We then categorized intratumoral CD8 T cells based on the median of the polyfunctionality score and compared the *Ythdf2* expression level between the high and low polyfunctionality groups. The resulting graphs were plotted using the R package ggplot2 (version 3.3.6). In the GSE206325 dataset, the cells were subjected to quality control, standardization, clustering, and dimensionality reduction, resulting in 34 cell clusters. Using hallmark genes, these clusters were further categorized into seven subpopulations: Terminal 1, Progenitor 2, Proliferating 3, Effector 4, Effector 5, Memory 6, and Cytotoxic 7. We examined the differences in YTHDF2 expression levels among the seven subpopulations or based on the computed polyfunctionality score.

### Statistical analysis

Histological analyses of both mouse and human tissue were performed in a blinded fashion. Immunoblot and immunofluorescence images are representative of experiments that have been repeated at least three times with similar results. Data from cell culture-based flow cytometry and qPCR assays were generated from two or three independent experiments. Data are presented as mean ± standard error of the mean (SEM). The statistical significance of differences was evaluated by two-tailed unpaired Student’s t test, two-sided Wilcoxon tests or one(two)-way ANOVA. *P* values of less than 0.05 were considered statistically significant. All statistical analyses were carried out using R (v4.2.2) or Graphpad Prism 8 (GraphPad Software).

### Study approval

All patient samples were obtained from Sun Yat-sen University Cancer Center (SYSUCC). The collection of tissue specimens was approved by the internal review and ethics boards of SYSUCC. All animal care and handling procedures were performed in accordance with the NIH’s Guide for the Care and Use of Laboratory Animals (National Academies Press, 2011) and were approved by the ethics committees of Sun Yat-sen University and University of Macau.

### Data availability

Data for bulk RNA-seq, RIP-seq, meRIP-seq, ATAC-seq, Ribo-seq and CUT&RUN are available through the BioProject portal (BioProject ID: PRJNA748842). All other data are available from the corresponding author on reasonable request.

## Acknowledgements

We thank Prof. Z. Yin (Jinan University) for providing the *Mettl3*^F/F^ mice. We thank Dr. R. Su (Beckman Research Institute of City of Hope), Dr. Z. Chen (Harvard University) and Dr. H. Huang (Sun Yat-sen University) and Dr. K. Wu (University of Macau) for helpful discussion. This work was supported by Guangdong Provincial Science Fund for Distinguished Young Scholars (2021B1515020007, to J.H.), General Program of National Natural Science Foundation of China (8271881 & 81871970, to J.H.), Macau Science and Technology Development Fund (FDCT) (0071/2023/RIA2, to J.H.), CAMS Innovation Fund for Medical Sciences (CIFMS) (2019-I2M-5-036, to R.-H. X.), Science and Technology Program of Guangdong (2019B020227002, to R.-H. X.), Hundred Talents Program of Sun Yat-sen University (2019079, to J.H.) and Ministry of Education Frontiers Science Centre for Precision Oncology, University of Macau (SP2023-00001-FSCPO, to J.H.).

## Author contributions

H.Z performed the animal and immunological experiments, interpreted the data and prepare the manuscript. X.L., Z.W., Zhicong Zhao, X.P., J.H.L., and W.W. performed the cellular and molecular experiments. W.Y., X.Z., and Q.S. performed the animal experiments. C.C., Q.Z. and Y.C. analysed the multiple sequencing data. M.Q., Zhaolei Zeng and M.C. collected the clinical samples and patient information. C.D., J.C., B.S., J.L. and H.-Q.J. provided key suggestions. R.-H.X. and J.H. co-supervised the study. J.H. conceived the project, designed the study, interpreted the data and wrote the manuscript. The order of the co–first authors was determined according to the time spent on this project.

## Competing interests

J.C. is a scientific founder of Genovel Biotech Corp. and holds equities with the company, and is also a Scientific Advisor for Race Oncology. Other authors declare no conflict of interests.

## Supplementary Figure Legends

**Supplementary Fig. 1.**
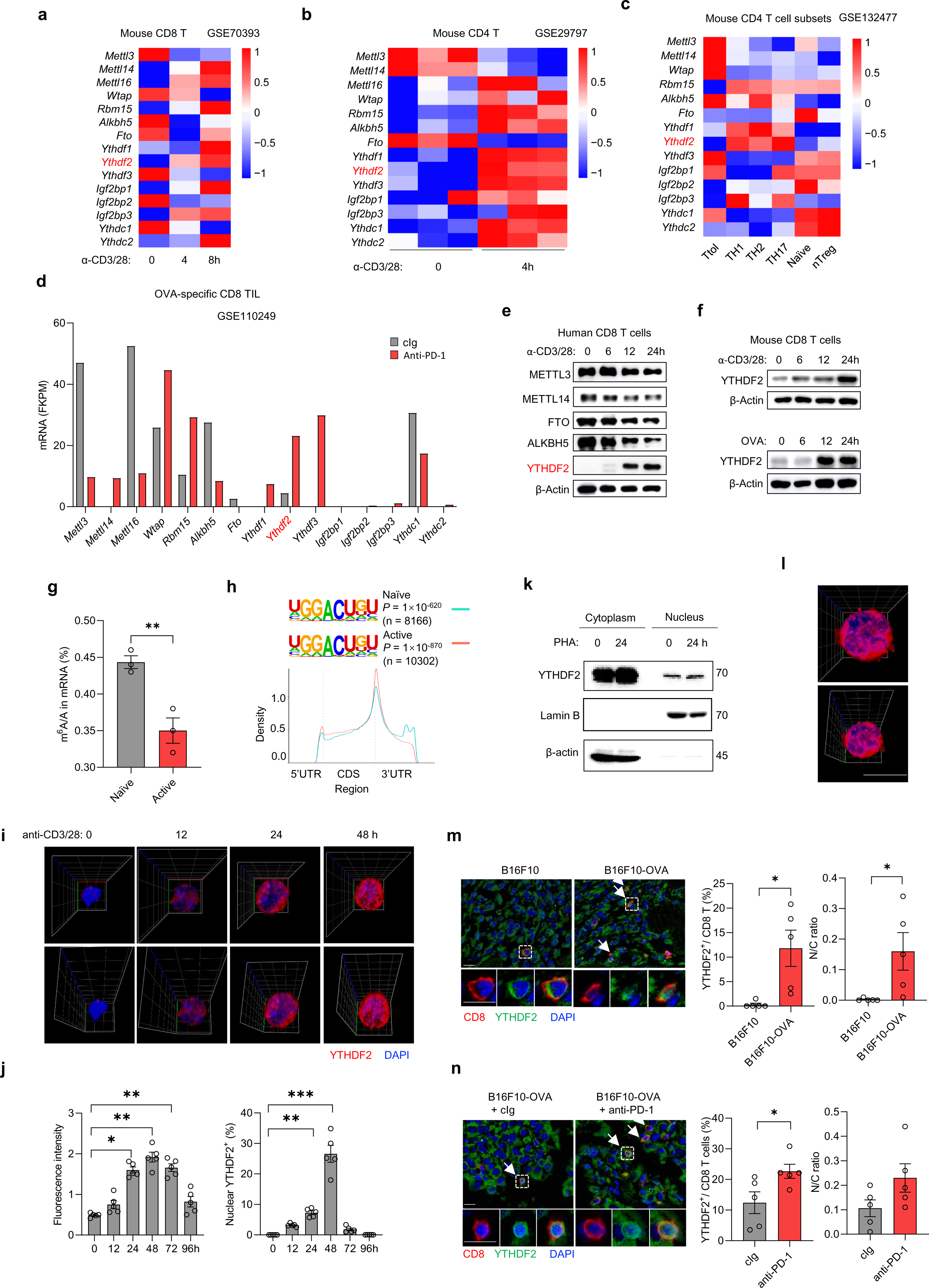
YTHDF2 is characteristically expressed upon T cell activation and reinvigoration. **a–d** Transcriptomic data were mined from existing datasets. mRNA expression of m^6^A modifiers in mouse naïve or activated WT CD8 (a) and CD4 (b) T cells. mRNA expression of m^6^A modifiers in T tolerant (T_tol_), T helper (T_H_1, T_H_2 and T_H_17), natural regulatory T (nT_reg_) and naïve T cells (c). mRNA expression of m^6^A modifiers in OVA-specific CD8 TILs from anti-PD-1- or cIg-treated mouse tumors (d). **e–f** Protein expression of m^6^A modifiers in CD8 T cells primed for the indicated amount of time. One representative of three independent experiments is shown. Human CD8 T cells from peripheral blood were stimulated with 5 μg/ml anti-CD3/CD28 (e). Wild-type (WT) (top) or OT-1 mouse CD8 T cells (bottom) were stimulated with 5 μg/ml anti-CD3/CD28 or 10 nM OVA (f). **g** LC-MS/MS-based m^6^A quantification in mRNAs from naïve and activated CD8 T cells (anti-CD3/CD28, 5 μg/ml, 24 h) (n = 3 per group). **h** Metagene distribution of the m^6^A peaks of naïve and active CD8 T cells (anti-CD3/CD28, 5 μg/ml, 24 h) along the whole transcriptome. The enriched consensus motifs were detected within m^6^A peaks. **i** Representative confocal Z-stack images of YTHDF2 (red), and DAPI (blue) in naïve or activated (anti-CD3/CD28, 5 μg/ml, 0–48 h) CD8 T cells. **j** Quantification of YTHDF2 intensity of CD8 T cells (left) and frequencies of nuclear YTHDF2^+^ cells from naïve or activated (anti-CD3/CD28, 5 μg/ml, 0–96 h) (right) CD8 T cells (n = 5 per group). **k** Immunoblotting analysis of YTHDF2 in the cytosol and nucleus of Jurkat cells stimulated with or without PHA (150 ng/ml, 24 h). respectively. Relative YTHDF2 levels were calculated using densitometry values for β-actin or Lamin B as calibrators. **l** Representative confocal Z-stack images of YTHDF2 (red) and DAPI (blue) in unstimulated Jurkat cells. Scale bar, 10 μm. **m** Representative multiplexable immunofluorescent staining of CD8 (red) and YTHDF2 (green) within B16F10 or B16F10-OVA tumors grown in OT-1 mice (n = 5 per group). A dashed box represents the 4× enlarged area shown in the bottom panels with separate channels. White arrows point to cells positive for YTHDF2 and CD8. Scale bar, 10 μm. Middle panel, frequencies of YTHDF2-positive CD8 T cells. Right panel, quantification of the nuclear to cytoplasmic ratios of YTHDF2 intensity in YTHDF2-positive CD8 T cells. **n** Representative multiplexable immunofluorescent staining of CD8 (red) and YTHDF2 (green) within B16F10-OVA tumors treated with cIg or anti-PD-1(n = 5 per group). A dashed box represents the 4× enlarged area shown in the bottom panels with separate channels. White arrows point to cells positive for YTHDF2 and CD8. Scale bar, 10 μm. Middle panel, frequencies of YTHDF2-positive CD8 T cells. Right panel, quantification of the nuclear to cytoplasmic ratios of YTHDF2 intensity in YTHDF2-positive CD8 T cells. Error bars, mean ± s.e.m. \**P* < 0.05; \*\**P* < 0.01; \*\*\**P* < 0.001. Two-tailed unpaired Student’s t-test (j, m, n).

**Supplementary Fig. 2.**
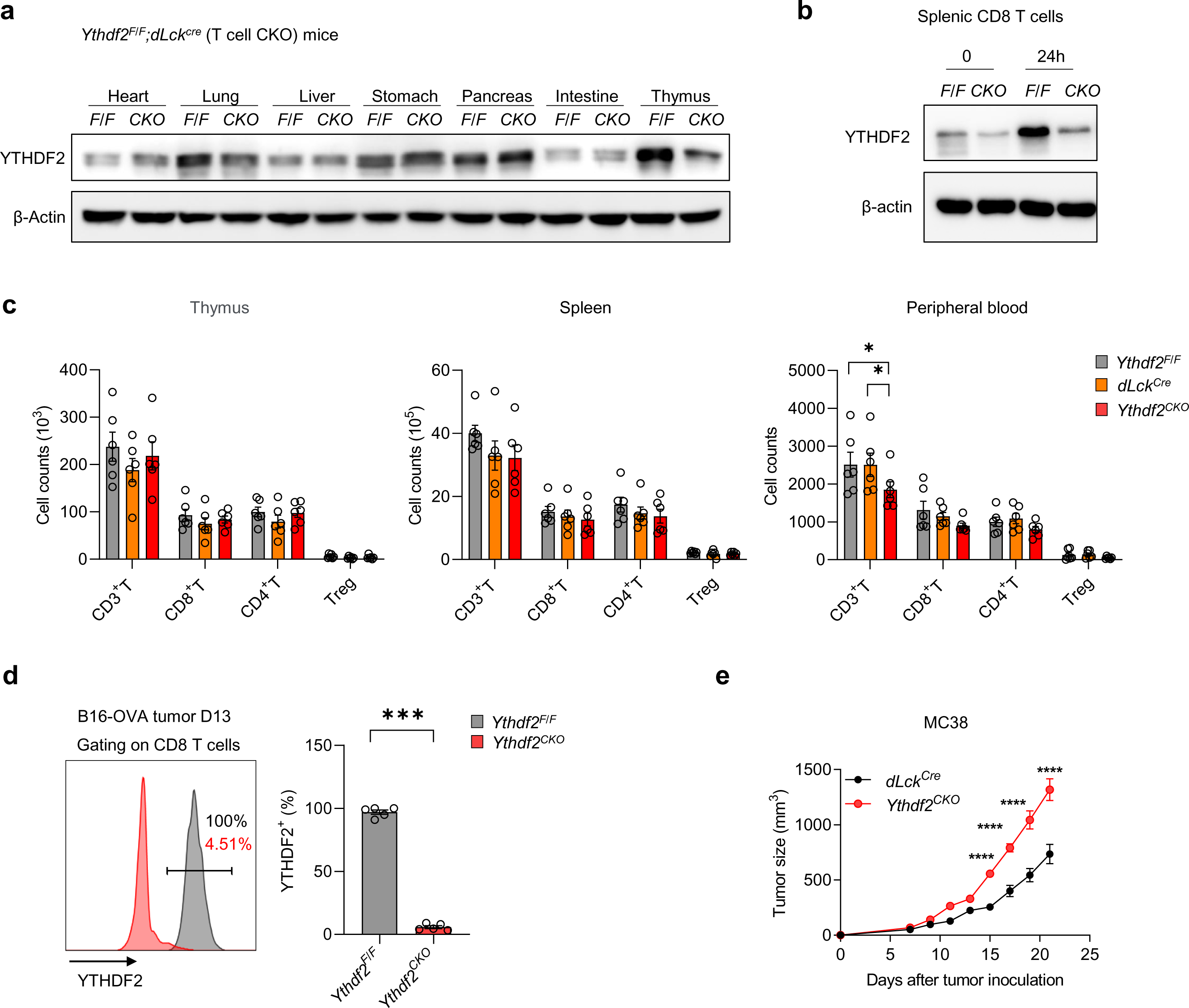
Phenotyping for the YTHDF2 conditional knockout miceYTHDF2 deprivation hinders antitumor T cell immune response. **a** Immunoblotting analysis of YTHDF2 expression in heart, lung, liver, stomach, pancreas, intestine and thymus tissues from *Ythdf2^F/F^* and *Ythdf2^CKO^* mice. **b** Protein expression of YTHDF2 in naïve and activated CD8 T cells (anti-CD3/CD28, 5 μg/ml, 24 h) derived from *Ythdf2^F/F^* and *Ythdf2^CKO^* mouse spleens. **c** Flow cytometry analysis of immune cells within the thymuses, spleens, peripheral blood from *Ythdf2^F/F^*, *dLck^Cre^* and *Ythdf2^CKO^* mice (n = 6 per group). **d** Quantification of YTHDF2 expression in B16-OVA tumor-infiltrating CD8 T cells from *Ythdf2^F/F^* and *Ythdf2^CKO^* mice (n = 5 per group). **e** Female *dLck^Cre^* (n = 5) and *Ythdf2^CKO^* (n = 5) mice were injected subcutaneously with 10^6^ MC38 cells. Tumor growth was monitored ever 2 or 3 days. Error bars, mean ± s.e.m. \**P* < 0.05; \*\*\**P* < 0.001; \*\*\*\**P* < 0.0001. One-way (c) or two-way ANOVA (e) or two-tailed unpaired Student’s t-test (d).

**Supplementary Fig. 3.**
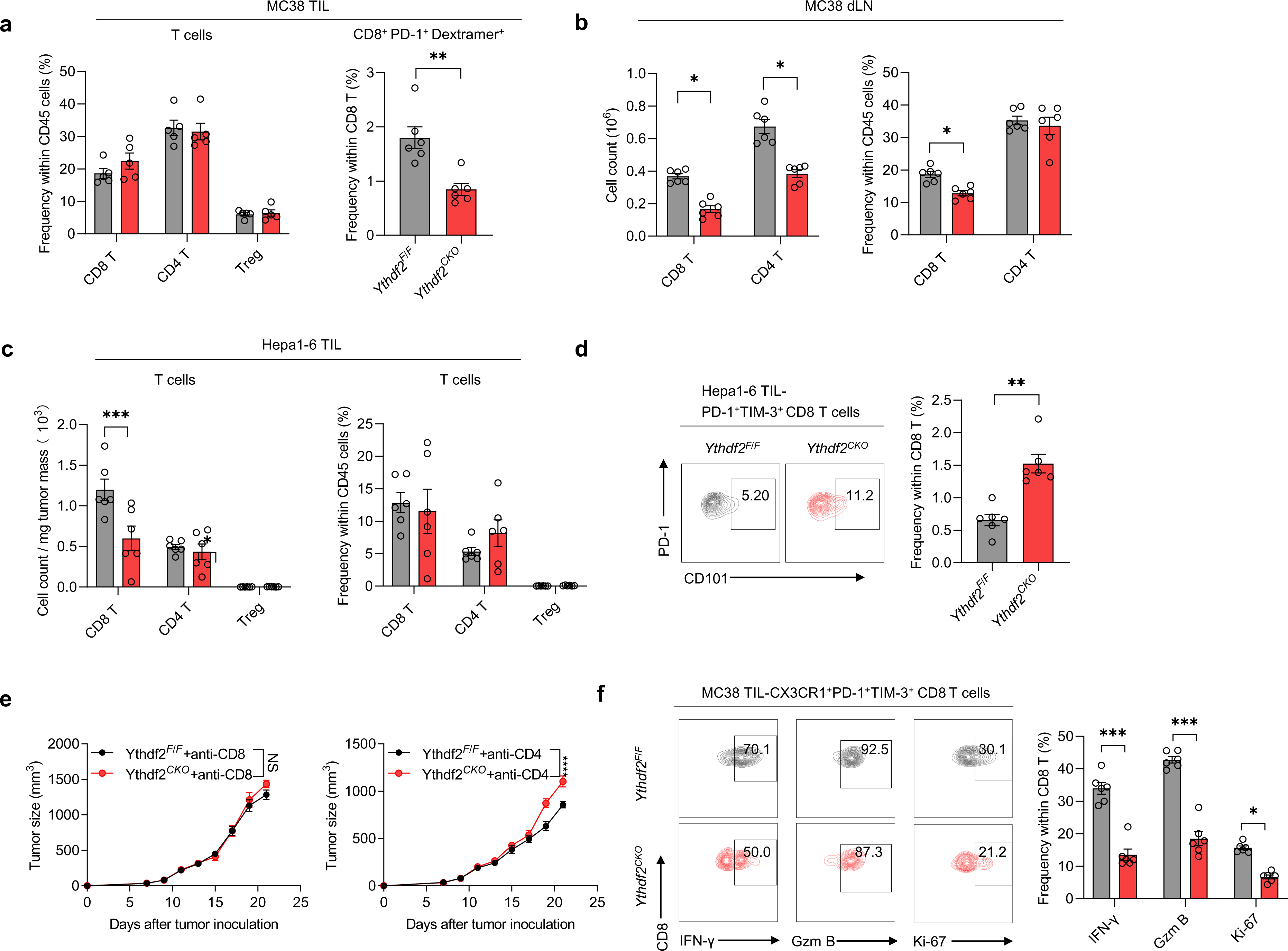
YTHDF2 deprivation hinders antitumor T cell immune response. **a** TILs were isolated from *Ythdf2^F/F^* (n = 5) and *Ythdf2^CKO^* (n = 5) mice 12 days after MC38 tumor inoculation. Frequencies of immune subsets (CD8 T, CD4 T and T_reg_ cells) within TILs and frequencies of PD-1^+^KILTFDRL-Dextramer^+^ CD8 T cell subpopulations were assessed by flow cytometry. **b** Numbers and frequencies of immune subsets within tumor-draining lymph nodes (dLN) from MC38 tumor-bearing *Ythdf2^F/F^* (n = 6) and *Ythdf2^CKO^* (n = 6) mice. **c** TILs were isolated from *Ythdf2^F/F^* (n = 6) and *Ythdf2^CKO^* (n = 6) mice 10 days after Hepa1-6 tumor inoculation. Frequencies of immune subsets (CD8 T, CD4 T and T_reg_ cells) within TILs and frequencies of PD-1^+^KILTFDRL-Dextramer^+^ CD8 T cell subpopulations were assessed by flow cytometry. **d** Frequencies of PD-1^+^TIM3^+^CD101^+^ CD8 T cell subpopulations from Hepa1-6 tumor-bearing *Ythdf2^F/F^* (n = 6) and *Ythdf2^CKO^* (n = 6) mice (Day 14). **e** MC38-bearing *Ythdf2^F/F^* (n = 5) and *Ythdf2^CKO^* (n = 5) mice were treated with anti-CD8 (200 μg/mouse) (left) or anti-CD4 antibody (200 μg/mouse) (right) and monitored for tumor growth. **f** Frequencies of CX3CR1^+^Tim3^+^CD101^-^ CD8 T cell subpopulations positive for Gzm B, IFN-γ or Ki-67 from MC38 tumor-bearing *Ythdf2^F/F^* (n = 6) and *Ythdf2^CKO^* (n =6) mice with anti-PD-1 treatment (D12). Error bars, mean ± s.e.m. \**P* < 0.05; \*\**P* < 0.01; \*\*\**P* < 0.001; \*\*\*\**P* < 0.0001. Two-way ANOVA (e) or two-tailed unpaired Student’s t-test (a–d, f).

**Supplementary Fig. 4.**
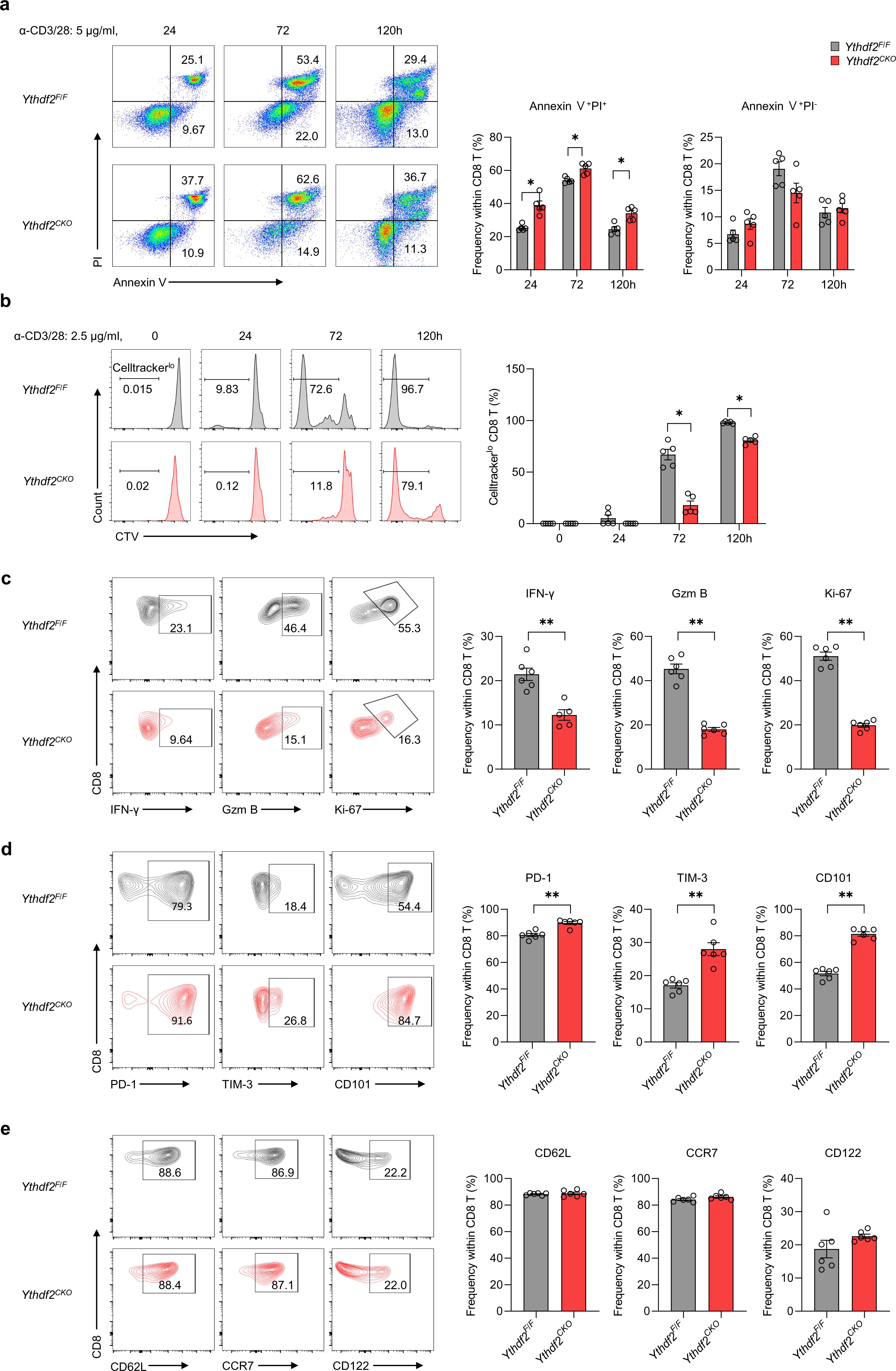
YTHDF2 maintains CD8 T cell expansion and activation *in vitro*. **a–c** Naïve CD8 T cells isolated from *Ythdf2^F/F^* (n = 5) or *Ythdf2^CKO^* (n = 5) mice were stimulated with anti-CD3/CD28 (2.5 or 5 μg/ml as indicated) for the indicated time. Apoptosis of *Ythdf2^F/F^* or *Ythdf2^CKO^* CD8 T cells was measured by Annexin V and propidium iodide (PI) staining (a). Proliferation of *Ythdf2^F/F^* or *Ythdf2^CKO^* CD8 T cells was assessed by Celltracker V (CTV) dilution (b). Quantification of frequencies of IFN-γ^+^, GZM B^+^, and Ki-67^+^ subpopulations within activated *Ythdf2^F/F^* or *Ythdf2^CKO^* CD8 T cells (c). **d** Naïve CD8 T cells isolated from *Ythdf2^F/F^* (n = 5) or *Ythdf2^CKO^* (n = 5) mice to induce T cell exhaustion by chronic stimulation (plates coated with anti-CD3, 5 mg/mL, 8 days). Quantification of frequencies of IFN-γ^+^, GZM B^+^, and Ki-67^+^ subpopulations within *Ythdf2^F/F^* or *Ythdf2^CKO^* CD8 T cells from in vitro T cell exhaustion assay. **e** Quantification of frequencies of CD62L^+^, CCR7^+^, and CD122^+^ subpopulations at Day 9 of transient stimulation (CD3/CD28 beads,1:1 beads-to-cells ratio, 72h) in *Ythdf2^F/F^* or *Ythdf2^CKO^* CD8 T cells from in vitro memory-like T cell induction assay (n = 6 per group). Error bars, mean ± s.e.m. \**P* < 0.05; \*\**P* < 0.01. Two-tailed unpaired Student’s t-test (a–e).

**Supplementary Fig. 5.**
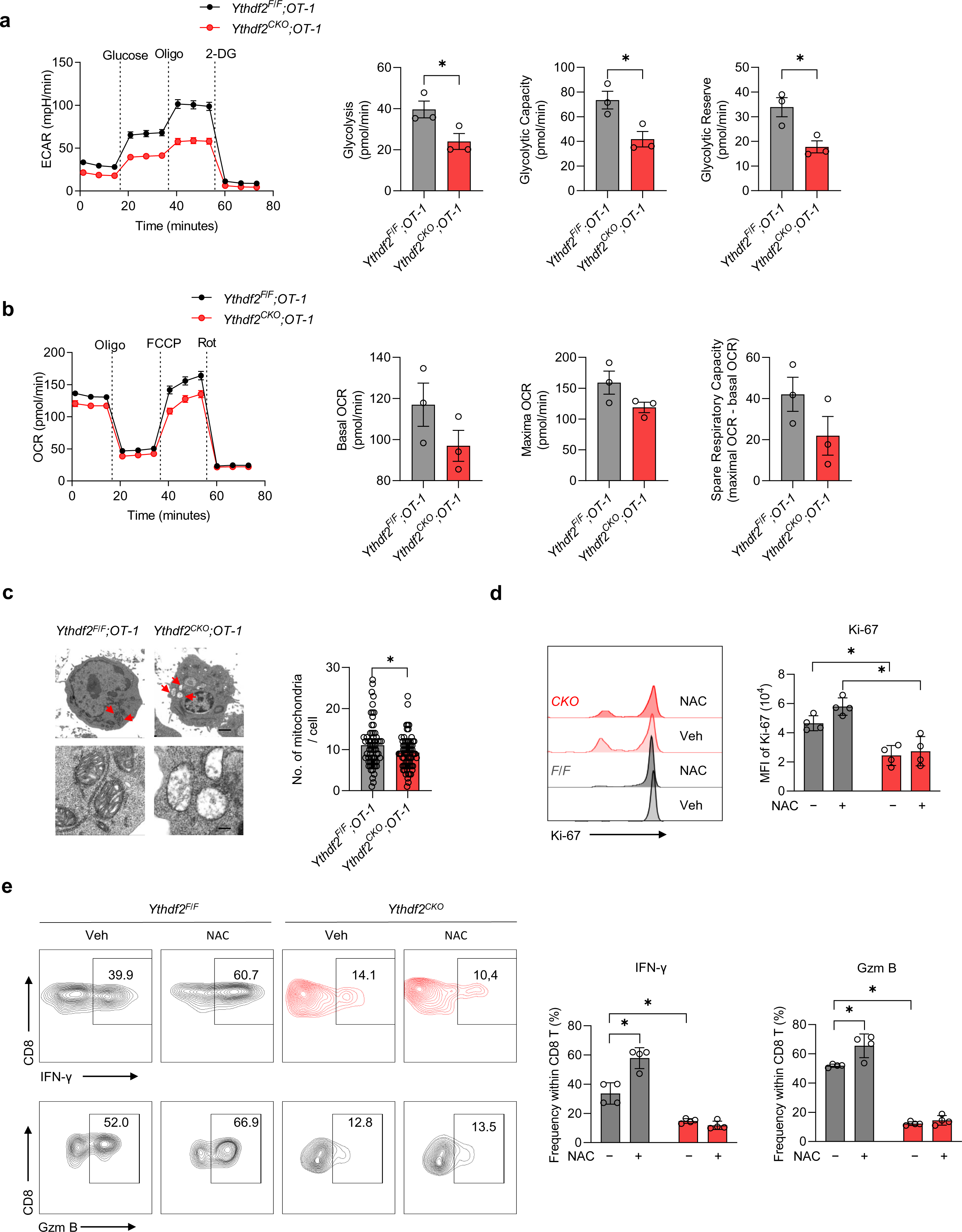
YTHDF2 deficiency impairs the mitochondria function of CD8 T cells. **a–b** Extracellular acidification rate (ECAR) levels (a) and oxygen consumption rate (OCAR) levels of primed *Ythdf2^F/F;^OT-1* or *Ythdf2^CKO^;OT-1* CD8 T cells (OVA, 10 nM, 72 h) were measured in real-time with Seahorse assay (n = 3 per group). **c** Transmission electron microscopic (TEM) analysis of mitochondrial morphology in activated *Ythdf2^F/F;^OT-1* or *Ythdf2^CKO^;OT-1* CD8 T cells (OVA, 10 nM, 72 h). Red arrows in the left panels show the position of mitochondria in respective higher magnification in the right panels. Scale bar, 200 nm. **d–e** Quantification of Ki-67 MFI (d), IFN-γ^+^, GZM B^+^ (e) frequencies among *Ythdf2^F/F^* and *Ythdf2^CKO^* CD8 T cells primed in the T cell exhaustion assay (plates coated with anti-CD3, 5 mg/mL, 8 days) with 10 mΜ NAC or veh (72 h) (n = 4 per group). Error bars, mean ± s.e.m. \**P* < 0.05. Two-tailed unpaired Student’s t-test (a–c). Two-way ANOVA (d, e).

**Supplementary Fig. 6.**
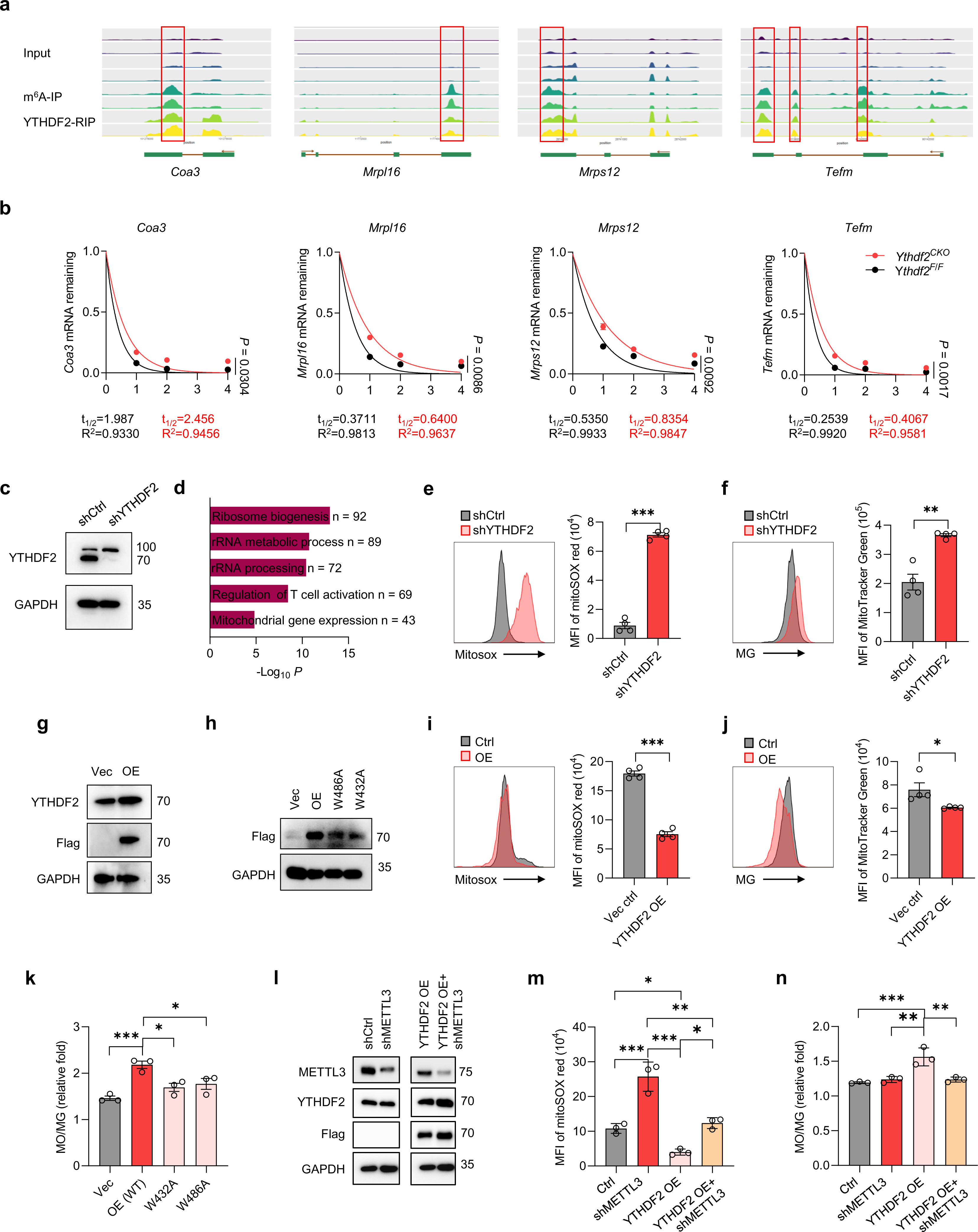
The m^6^A machinery is necessary for YTHDF2-regulated mitochondrial fitness. **a** m^6^A-seq and RIP-seq tracks of *Coa3, Mrpl16, Mrps12* and *Tefm* mRNA loci on primed*Ythdf2^F/F^* and *Ythdf2^CKO^* CD8 T cells. **b** Activated *Ythdf2^F/F^* and *Ythdf2^CKO^* CD8 T cells (anti-CD3/CD28, 5 μg/ml, 24 h) were treated with ActD (500 μg/ml) and RNAs were collected at different time points. *Coa3, Mrpl16, Mrps12 and Tefm* mRNA levels were measured using qPCR and represented as mRNA remaining after ActD treatment (n = 3 per group). **c** Immunoblotting analysis of YTHDF2 expression in Jurkat-shCtrl and Jurkat-shYTHDF2 cells. **d** GO enrichment analysis of upregulated genes in Jurkat-shCtrl and Jurkat-shYTHDF2 cells. **e–f** Quantification of MitoSOX (e) and MG (f) MFI in Jurkat-shCtrl and Jurkat-shYTHDF2 cells (n = 4 per group). **g** Immunoblotting analysis of YTHDF2 and Flag expression in Jurkat cells with or without YTHDF2 overexpression (OE) (n = 4 per group). **h** Immunoblotting analysis of Flag expression in Jurkat cells transduced with wild-type (OE) or mutant YTHDF2 (W432A and W486A) or empty vector lentiviruses. **i–j** Quantification of MitoSOX (i) and MG (j) MFI in Jurkat cells with or without YTHDF2 OE (n = 4 per group). **k** Quantification of the MO/MG ratio in Jurkat cells transduced with wild-type or mutant YTHDF2 or empty vector lentiviruses (n = 3 per group). **l** Immunoblotting analysis of METTL3, YTHDF2 and Flag expression in YTHDF2-OE Jurkat cells with or without knockdown of METTL3. **m** MitoSOX staining among METTL3-knockdown and control Jurkat cells with or without YTHDF2 overexpression (n = 3 per group). **n** Quantification of the MO/MG ratio among METTL3-knockdown and control Jurkat cells with or without YTHDF2 overexpression (n = 3 per group). Error bars, mean ± s.e.m. \**P* < 0.05; \*\**P* < 0.01; \*\*\**P* < 0.001. Two-tailed unpaired Student’s t-test (f, i, j) or one-way ANOVA (k, m, n) or non-linear regression (b).

**Supplementary Fig. 7.**
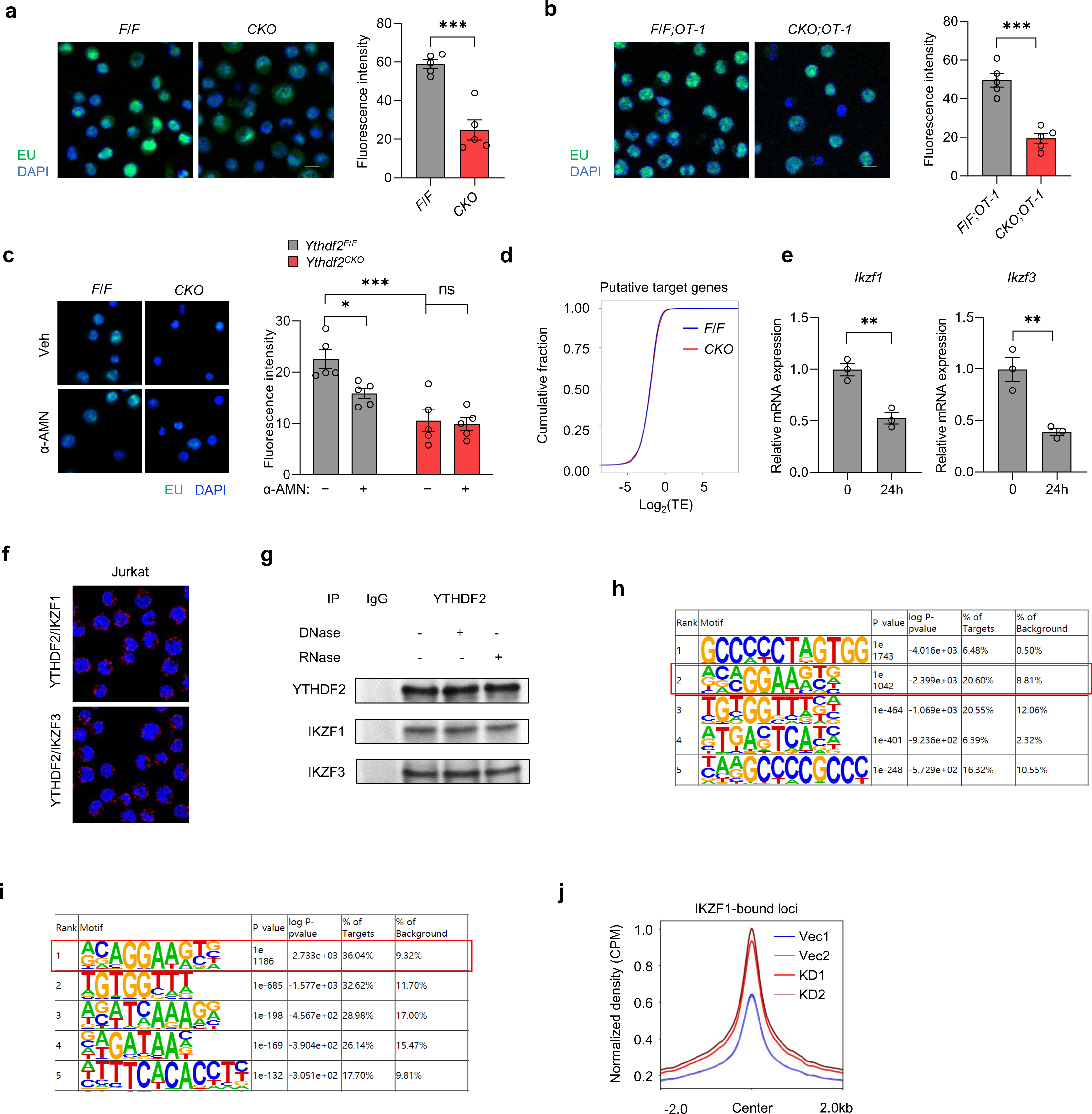
Acute T cell activation enlists nuclear functionality of YTHDF2. **a–b** Click-it RNA imaging and analysis of nascent RNA (green) synthesis in activated *Ythdf2^F/F^* or *Ythdf2^CKO^* CD8 T cells (anti-CD3/CD28, 5 μg/ml, 24 h) (a) and activated *Ythdf2^F/F;^OT-1* or *Ythdf2^CKO^;OT-1* CD8 T cells (OVA, 10 nM, 24 h) (b). Scale bar, 10 μm. **c** Click-it RNA imaging and analysis of nascent RNA (green) synthesis in activated *Ythdf2^F/F^* or *Ythdf2^CKO^* CD8 T cells (anti-CD3/CD28, 5 μg/ml, 24 h) in the presence of α-amanitin (2μg/mL, 12 h) or vehicle (n = 5 per group). Scale bar, 10 μm. **d** Cumulative distribution of the fold change in translational efficiency of YTHDF2-targeted and m^6^A-marked transcripts between activated *Ythdf2^F/F^* and *Ythdf2^CKO^* CD8 T cells (anti-CD3/CD28, 5 μg/ml, 24 h). **e** *Ikzf1 and Ikzf3* mRNA levels detected by qPCR in naïve or activated (anti-CD3/CD28, 5 μg/ml, 24 h) CD8 T cells (n = 3 per group). **f** PLA analysis of YTHDF2 associated with IKZF1 or IKZF3 in unstimulated Jurkat cells. Scale bar, 10 μm. **g** Whole-cell lysates of activated WT CD8 T cells (anti-CD3/CD28, 5 μg/ml, 24 h) were subjected to immunoprecipitation using anti-YTHDF2 antibody. The immunoprecipitants were incubated with RNase A (1 ug/ul) or DNase I (0.4 U/ul) DNase followed by immunoblot analysis. **h–i** Motif discovery analysis of the genomic sequences under ATAC-seq peaks conditioned by YTHDF2 depletion using HOMER. Red rectangle shows the IKZF1/3-binding motif within ATAC-sequenced *Ythdf2^F/F^* and *Ythdf2^CKO^* CD8 T cells (h) or Jurkat-shCtrl and Jurkat-shYTHDF2 cells (i). **j** ChIP-seq datasets for IKZF1 (GSM935442) in human T cells were obtained using Cistrome Data Browser. ATAC-seq profiles of Jurkat-shCtrl and Jurkat-shYthdf2 cells were represented on IKZF1-bound loci. Error bars, mean ± s.e.m. \**P* < 0.05; \*\**P* < 0.01; \*\*\**P* < 0.001. Two-tailed unpaired Student’s t-test (a, b, e) or two-way ANOVA (c).

**Supplementary Fig. 8.**
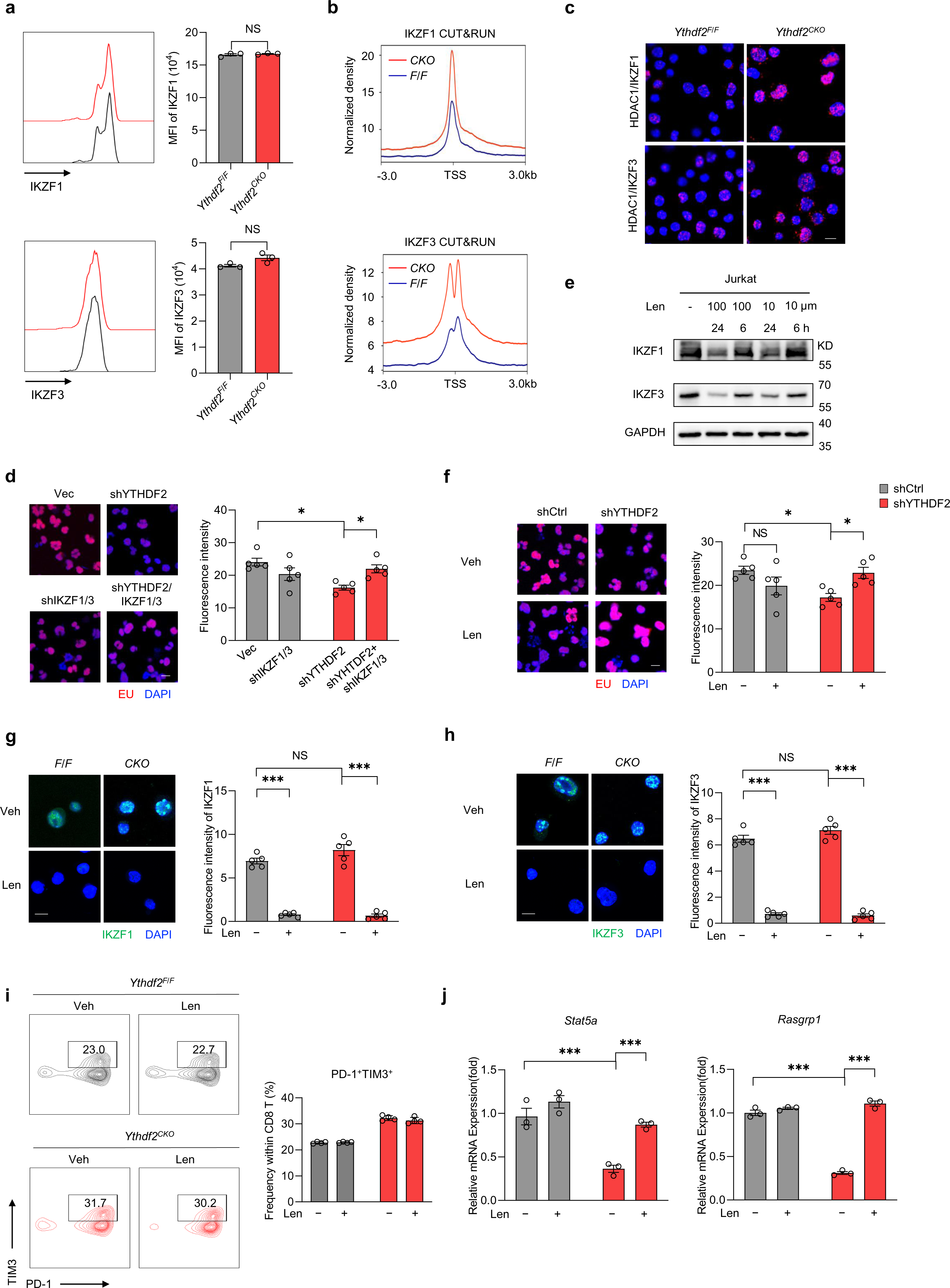
IKZF1/3-associated transcriptional repression in YTHDF2-deficient T cells. **a** Quantitative flow cytometry analysis of IKZF1 (top) or IKZF3 (bottom) expression in activated *Ythdf2^F/F^* and *Ythdf2^CKO^* CD8 T cells (anti-CD3/CD28, 5 μg/ml, 24 h) (n = 3). **b** IKZF1 (top) or IKZF3 (bottom) CUT&RUN profiles of activated *Ythdf2^F/F^* and *Ythdf2^CKO^* CD8 T cells (anti-CD3/CD28, 5 μg/ml, 24 h) were represented at the gene promoter regions. **c** PLA analysis of HDAC1 associated with IKZF1 or IKZF3 in activated *Ythdf2^F/F^* and *Ythdf2^CKO^* CD8 T cells (anti-CD3/CD28, 5 μg/ml, 24 h). Scale bar, 10 μm. **d** Click-it RNA imaging and analysis of nascent RNA synthesis (red) in Jurkat-shCtrl, Jurkat-shIKZF1/3 cells, Jurkat-shYTHDF2 cells and Jurkat-shYTHDF2/IKZF1/3 cells (n = 5 per group). **e** Immunoblotting analyses of IKZF1 and IKZF3 in Jurkat cells treated with len (10 or 100 μΜ) or veh for 6 or 24 h. **f** Click-it RNA imaging and analysis of nascent RNA synthesis (red) in Jurkat-shCtrl and Jurkat-shYTHDF2 cells in the presence of 100 μΜ len or veh for 24 h (n = 5 per group). **g**–**h** Representative confocal immunofluorescence images of IKZF1 (g) or IKZF3 (h) (green) and DAPI (blue) in activated *Ythdf2^F/F^* and *Ythdf2^CKO^* CD8 T cells (anti-CD3/CD28, 5 μg/ml, 24 h) in the presence of 10 μΜ len or veh (n = 5 per group). Scale bar, 10 μm. **i** Quantification of Tim3^+^ PD-1^+^ frequencies among primed*Ythdf2^F/F^* and *Ythdf2^CKO^* CD8 T cells (anti-CD3/CD28, 5 μg/ml, 48 h) in the presence of 10 μΜ len or veh (n = 4 per group). **j** *Stat5a* (left) and *Rasgrp1* (right) mRNA levels detected by qPCR in activated *Ythdf2^F/F^* and *Ythdf2^CKO^* CD8 T cells (anti-CD3/CD28, 5 μg/ml, 24 h) in the presence of 10 μΜ len or veh (n = 3 per group). Error bars, mean ± s.e.m. \**P* < 0.05; \*\**P* < 0.01; \*\*\**P* < 0.001. Two-tailed unpaired Student’s t-test (a). One (d) or two-way (f–j) ANOVA.

**Supplementary Fig. 9.**
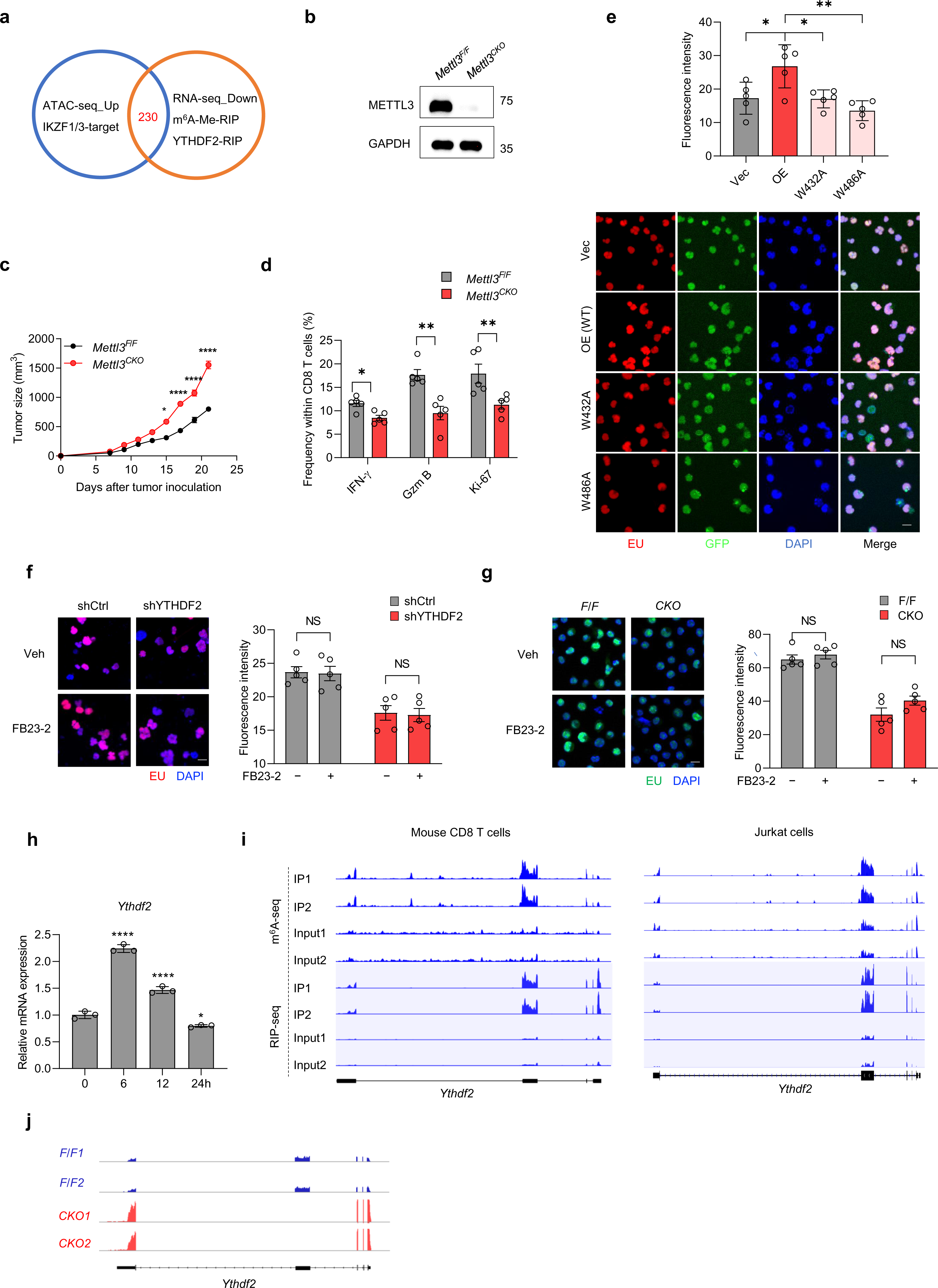
YTHDF2 distribution and expression are governed by the m6A machinery. **a** Venn diagram showing the overlap of putative YTHDF2-target genes and IKZF1/3-target genes with both enhanced chromatin accessibility and reduced mRNA expression found in *Ythdf2^CKO^* CD8 T cells. **b** Immunoblotting analysis of METTL3 and GAPDH expression in activated *Mettl3^F/F^* or *Mettl3^CKO^* CD8 T cells (anti-CD3/CD28, 5 μg/ml, 24 h). **c** Female *Mettl3^F/F^* (n = 6) and *Mettl3^CKO^* (n = 6) mice were injected subcutaneously with 10^6^ MC38 cells. Tumor growth was monitored ever 2 or 3 days. **d** TILs were isolated from *Mettl3^F/F^* (n = 5) and *Mettl3^CKO^* (n = 5) mice 12 days after MC38 tumor inoculation. Frequencies of CD8 T cell subpopulations positive for Gzm B, IFN-γ, or Ki-67 were assessed by flow cytometry. **e** Click-it RNA imaging and analysis of nascent RNA (red) synthesis in Jurkat cells introduced with WT or mutant YTHDF2. Scale bar, 10 μm (n = 5 per group). **f** Click-it RNA imaging and analysis of nascent RNA synthesis (red) in Jurkat-shCtrl and Jurkat-shYTHDF2 cells in the presence of 10 μΜ FB23-2 or veh for 72 h (n = 5 per group). Scale bar, 10 μm. **g** Click-it RNA imaging and analysis of nascent RNA synthesis (green) in *Ythdf2^F/F^* and *Ythdf2^CKO^* CD8 T cells primed in the presence of 10 μΜ FB23-2 or veh for 72 h (n = 5 per group). Scale bar, 10 μm. **h** *Ythdf2* mRNA levels detected by qPCR in CD8 T cells stimulated with anti-CD3/CD28 (5 μg/ml) for the indicated amount of time (n = 3 per group). **i** m^6^A-seq and RIP-seq tracks of *Ythdf2* mRNA loci on mouse CD8 T cells (left) and Jurkat cells (right). **j** RNA-seq tracks of *Ythdf2* mRNA loci on primed*Ythdf2^F/F^* and *Ythdf2^CKO^* CD8 T cells. Error bars, mean ± s.e.m. NS, no significance; \**P* < 0.05; \*\*\*\**P* < 0.0001. Two-tailed unpaired Student’s t-test (d). One-way (e, h) or two-way ANOVA (c, f, g).

**Supplementary Fig. 10.**
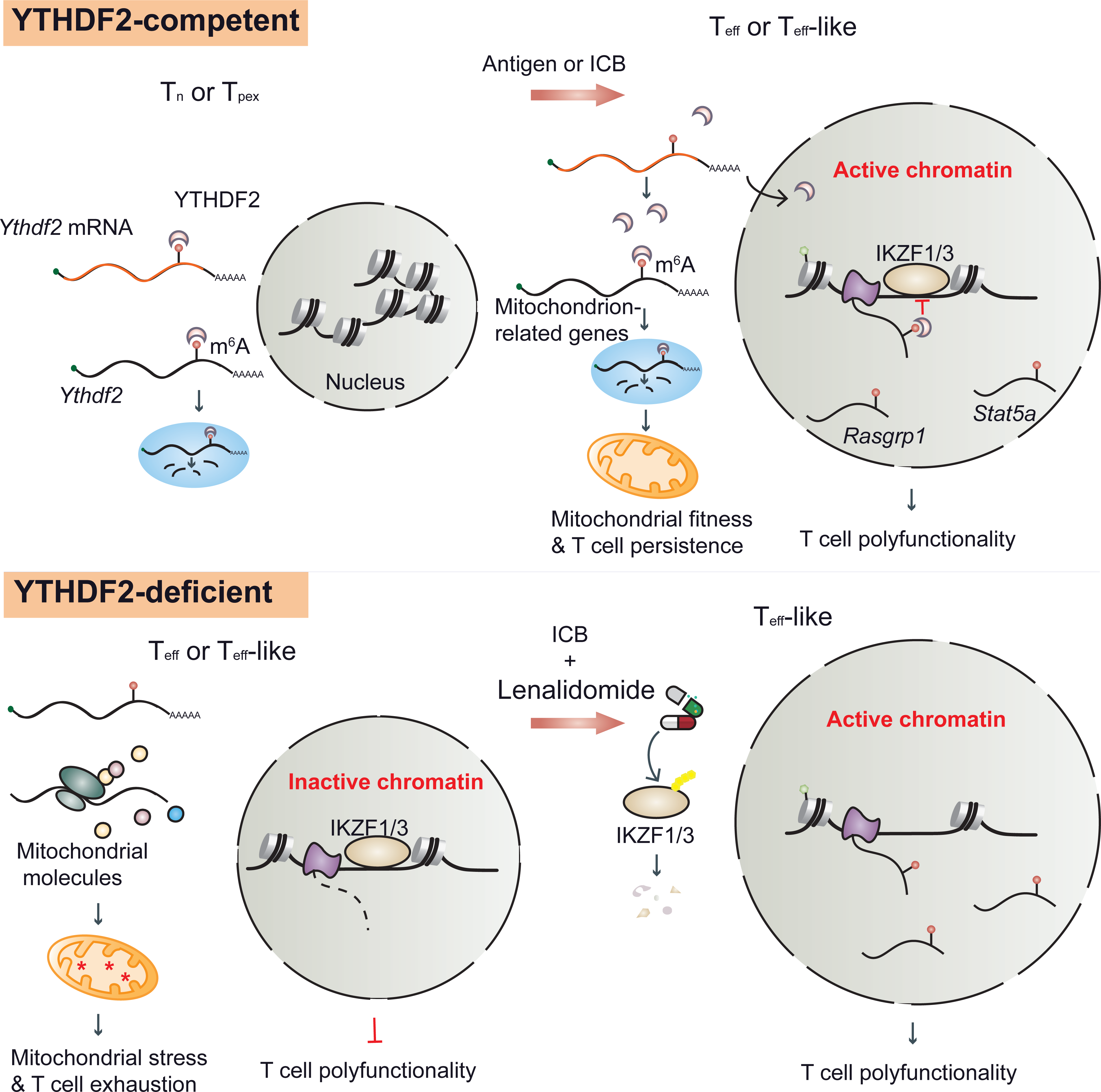
Schematic diagram of YTHDF2 functioning in antitumor CD8 T cells. Contrary to the autoregulated *Ythdf2* mRNA decay in a quiescent state, YTHDF2 protein is partially relocated and swiftly accumulated upon early CD8 T cell activation or reinvigoration. While cytoplasmic YTHDF2 degrades redundant mitochondrial component-encoding mRNAs to sustain T cell persistence, its nuclear translocation is likely to safeguard T cell effectiveness and ICB responsiveness by minimizing IKZF1/3-mediated transcriptional repression. Conversely, YTHDF2 defect-associated ICB resistance could be overcome by targeting IKZF1/3.

